# Chemotherapy coupled to macrophage inhibition leads to enhanced T and B cell infiltration and durable triple negative breast cancer regression

**DOI:** 10.1101/2021.02.22.432300

**Authors:** Swarnima Singh, Nigel Lee, Diego A. Pedroza, Igor Bado, Licheng Zhang, Sergio Aguirre, Clark Hamor, Yichao Shen, Yitian Xu, Jingyuan Hu, Yang Gao, Na Zhao, Shu-Hsia Chen, Ying-Wooi Wan, Zhandong Liu, Jeffrey T. Chang, Daniel Hollern, Charles M. Perou, Xiang H.F. Zhang, Jeffrey M. Rosen

## Abstract

Immunosuppressive elements within the tumor microenvironment such as Tumor Associated Macrophages (TAMs) can present a barrier to successful anti-tumor responses by cytolytic T cells. We employed preclinical syngeneic p53 null mouse models of triple negative breast cancer (TNBC) to develop a treatment regimen that harnessed the immunostimulatory effects of low-dose cyclophosphamide coupled with the pharmacologic inhibition of TAMs using either a small molecule CSF1R inhibitor or an anti-CSF1R antibody. This therapeutic combination was used to successfully treat several highly aggressive TNBC murine mammary tumors and lung metastasis. Using this regimen and single cell RNA sequencing we characterized tumor infiltrating lymphocytes (TILs) including helper T cells and antigen-presenting B cells that were highly enriched in good responders to combination therapy. Using high dimensional imaging techniques, we identified the close spatial localization of B220+ CD86+ activated B cells and CD4+ T cells in tertiary lymphoid structures that were present up to 6 weeks post-treatment in one model that also exhibited long-term tumor regression post-treatment. We also characterized the transcriptional and metabolic heterogeneity of TAMS in these two closely related claudin-low/mesenchymal subtype tumor models with differential treatment responses. A murine TAM signature derived from the T12 model is highly expressed and conserved in human claudin-low breast cancers, and high expression of the T12 signature correlated with reduced overall survival. This T12 tumor TAM signature may help identify human claudin-low breast cancer patients that will benefit from the combination of cyclophosphamide and anti-CSF1R therapy. These studies illustrate the complexity of the tumor immune microenvironment and highlight different immune responses that result from rationale combinations of immunotherapy.

**Significance:** A treatment regimen that harnessed the immunostimulatory effects of cyclophosphamide coupled with the inhibition of CSF1R was used to successfully treat several highly aggressive claudin-low TNBC murine mammary tumors and lung metastasis.

## Introduction

Triple-negative breast cancer (TNBC) is a heterogeneous group of breast cancers defined by the absence of ER, PR and Her2. TNBC disproportionately affects young women and especially those of African ancestry and is an aggressive subtype of breast cancer with an overall poorer prognosis compared to other breast cancer subtypes (1). At present, the primary systemic treatment option for TNBC in the adjuvant setting is multi-agent chemotherapy. Many TNBCs are chemotherapy sensitive, and patients have pathological complete response (pCR) rates of 30–53% when treated with an anthracycline/taxane containing regimen(1–3). However, those patients who do not achieve a pCR have a poor prognosis (4). Immunotherapy is now approved for use in PD-L1^+^ metastatic TNBC patients as well as neoadjuvant treatment of TNBC; but even among PD-L1^+^ patients the response is variable(5). Promising results in TNBC patients treated with immune checkpoint inhibitors and chemotherapy have been reported where the pCR rate was 65% compared to 51% with chemotherapy alone(6). Therefore, immune checkpoint inhibitors may play a role in the early treatment for TNBC patients, but the variability in therapeutic responses and patient outcomes remains a concern.

A common feature of TNBCs is their epithelial to mesenchymal transition (EMT) phenotype. EMT is an evolutionarily conserved developmental program during which cells lose epithelial markers and gain mesenchymal traits. EMT confers metastatic properties to cancer cells by enhancing mobility, invasion, and resistance to apoptotic stimuli. Moreover, intermediate or “partial EMT” tumor cells acquire increased plasticity and stemness properties, and exhibit marked therapeutic resistance(7). Our group reported the first results of neoadjuvant clinical trials in which residual breast cancers after conventional endocrine therapy (letrozole) or chemotherapy (docetaxel) displayed these intermediate EMT features and tumor-initiating properties(8).

We have developed multiple transplantable preclinical syngeneic TNBC genetically engineered mouse (GEM) models, which have been characterized genomically(9, 10) and with respect to their immune microenvironments(11, 12). By integrating the immunological characterization of murine syngeneic mammary tumor models with analyses of human breast cancer datasets, we have demonstrated a relationship between EMT and myeloid cells, specifically tumor associated macrophages (TAMs)(12). We also have leveraged our syngeneic GEM models to define the response to immune checkpoint blockade therapy (ICBT) with emphasis on the myeloid cell environment(12). Using these models, we have identified a role for neoantigens and the importance of B cells and T follicular helper cells(11). The EMT hallmark gene signature also has been found to be enriched in residual tumors after neoadjuvant chemotherapy in TNBC, and these tumors displayed a high level of residual polarized macrophages that can suppress T cell proliferation and activation (12). Claudin-low tumors, a subtype of TNBC, adopt a spindle-like morphology indicative of their highly mesenchymal nature. These TNBC claudin-low models have been characterized with respect to their immune microenvironment and have an enrichment in TAMs as compared to TNBC basal-like murine tumors. Furthermore, increasing the mutation burden in certain claudin-low tumors has been shown to improve their responsiveness to immunotherapy. These are, therefore, appropriate preclinical models in which to determine the functional importance of TAMs.

Single-cell RNA sequencing (scRNA-seq) as well as profiling of TCR repertoires have proven to be valuable tools to dissect the cellular heterogeneity in various cancers. By combining high dimensional transcriptomic data of individual tumor infiltrating immune cells with bulk RNA sequencing data from multiple murine mammary tumor models, we began to understand the complex dynamics that prevent tumors from undergoing long-term regression. Furthermore, we have harnessed the immunostimulatory effects of the low-dose chemotherapeutic agent cyclophosphamide (Cytoxan, CTX) aided by the depletion of immunosuppressive TAMs using either a small molecule inhibitor or monoclonal antibody towards CSF1R to successfully induce durable long-term responses in highly aggressive primary murine mammary tumors. Furthermore, we observed the presence of tertiary lymphoid structures (TLSs), identified B cell sub-populations that are enriched in responders and identified a macrophage signature that could be used to identify patients with Claudin-low breast cancer that might benefit from this treatment combination.

## Methods

### Cell Lines

T11 and T12 cell lines were generated in the Rosen laboratory from primary T11 and T12 tumors by selection of epithelial cells expressing the neomycin cassette in G418.and cultured in DMEM/High Glucose with 10 % FBS, 100 IU ml^-1^ penicillin/streptomycin (Lonza), 5 μg/ml insulin, 10 ng/ml mouse EGF, and AA) and hydrocortisone. Cells were maintained in a humidified incubator at 37°C with 5% CO2. Mycoplasma testing was carried out on the cell lines once a month.

### Animals Studies

All animals were used according to a protocol approved by the Institutional Animal Care and Use Committee at Baylor College of Medicine. Female 5-6-week-old BALB/c mice were ordered from Envigo Laboratories and experiments were carried out in age matched animals. Female 5-6-week-old NSG mice (NOD.Cg-Prkdc scid il2rg tm1 Wjl/SzJ) (Stock no 005557) were ordered from Jackson Labs. Mice were house in the TMF Mouse Facility at the Baylor College of Medicine in SPF/climate-controlled conditions with 12-hour day or night cycles. They were supplied with fresh chow and water from an autowater system continuously.

### Mammary fat pad injection

The generation of T12, T11 and 2151R tumor models has been described and characterized previouslyand tumor chunks were stored in liquid nitrogen for long-term storage in FBS + 10% DMSO. Prior to surgery, tumor chunks were thawed and washed in PBS. Tumor chunks were implanted directly into a small cavity in the mammary fat pad. Mice were monitored for tumor growth and were randomized into treatment groups when the tumors were approximately 5 mm in diameter.

### In-vivo treatment studies using Pexidartinib

Cyclophosphamide (Sigma Aldrich, PHR1404-1G) was resuspended in sterile PBS(Lonza) and injected i.p at the concentration of 100 mg/kg, once a week for the entire treatment period and the control mice were injected with the same volume of sterile PBS. PLX3397 was obtained from Plexxikon, Daichii Sankyo and was added to a chow made by Research Diets. Mice were fed the chow ad libitum. The drug concentration in the chow was 275 ppm in most of the experiments described which corresponds to 5 mg/kg per mouse except for the low-dose experiments with a concentration of 75 ppm. Control mice were given the same chow without PLX3397. Mice were weighed and monitored for signs of drug toxicity weekly. Tumors were measured using calipers 3 times a week for the duration of the treatment.

### In-vivo treatment studies using Axatilimab

For the studies using a high affinity anti-CSF1R antibody, Axatilimab (SNDX-ms6352) was obtained from Syndax Pharmaceuticals, Inc. and was administered via i.p injection weekly for four weeks at the concentration of 40mg/kg for the first dose followed by 20mg/kg for the remaining 3 doses. Control mice were injected with equal volume of mouse IgG1 isotype control (Bio X Cell, BE0083).

### In-vivo T cell depletion studies

Mice were injected i.p. with anti CD4(100 ug, BE0003-1) or anti CD8(150 ug, BE0061) antibodies obtained from Bioxcell Laboratories, twice times weekly. Optimal antibody concentrations were obtained from previous validation experiments. This was repeated during the entire treatment course.

### Tail vein injection

Female 5-7-week-old BALB/c mice were injected with tumor cells dissociated from fresh T12 mammary tumors. Cells (200,000) in sterile PBS were injected in per mouse using a 27-gauge syringe. Mice were randomized into treatment groups at day 10 post injection and were given 3 weekly treatments of CTX and PLX3397 chow ad libitum, following which the mice were sacrificed and the lungs were collected for further analysis.

### T cell isolation and In-vitro immunosuppression assay

Spleens were harvested from 8-week-old JEDI mice and physically dissociated using a scalpel. Splenocytes were filtered through a 70-m cell strainer. Red blood cells were lysed using RBC lysis buffer (#B4300062518TN, Tonbo Biosciences). CD3+ T cells were enriched using negative selection of biotinylated antibodies B220, CD11b, CD11c, and Gr-1 (#559971, BD Pharmingen) and magnetically sorting using EasySep Mouse Biotin Positive Selection kit (#18559, Stemcell). CD3+ T cells were activated using a 1:1 ratio Dynabead Mouse T Activator CD3/CD28 (#00775477, Thermo Scientific) for 72 hrs in T cell media containing RPMI 1640 (VWR), 5% heat inactivated FBS, 55 m ß-mercaptoethanol, and 5ng/ml IL-2(# 202-IL-050/CF, R&D Systems). Activated CD3+ T cells were isolated from Dynabead Mouse T Activator CD3/CD28 (#00775477, Thermo Fisher) using negative selection. T12 unlabeled and GFP labeled cell lines were seeded at a density of 2500 cells per well in a 96-well plate. After 24 hrs, T cells were added at a 10:1 effector/target ratio with 10,000; 1,000; 100; 10; 1; 0 m of phosphoramide mustard (#M123069, MuseChem). Each condition was run in triplicate. The conditions include co-culturing with T cells and drug, and wells with drug alone. The plate was incubated in an Incucyte (Essen) and fluorescence and phase images were taken every hour for 48 hrs. The analysis was run using Incucyte S3 software and graphed using Graphpad Prism 8. For co-culture experiments TAMs were isolated from fresh T12 or T11 tumors using the marker F480 by FACS sorting. They were seeded at a 1:1 ratio with the T cells in the media mentioned above with 20 % tumor cell conditioned medium. The cells were kept in culture for 72 hours following which the T cells and tumor cells were separated and analyzed by flow.

### Tissue staining

Primary tumor and lung tissue were fixed in 4% paraformaldehyde overnight and transferred to 70% ethanol for long-term storage. The tissue was embedded in paraffin blocks by the Breast Center Pathology Core at Baylor College of Medicine. IHC was performed according to Rosen Lab protocols (https://www.bcm.edu/research/labs/jeffrey-rosen/protocols). Primary antibodies used were ordered from Abcam-anti-CD4(EPR19514), Anti-CD8a (YTS169.4), S100A8(EPR3554), F480(CI: A3-1) or eBioscience B220(RA3-6B2). Biotinylated secondary antibodies were ordered from Biolegend.

### Tissue Quantification

Immunohistochemistry slides were quantified utilizing ImageJ. All images were first set to 8-bit black and white the same threshold was then applied to all images to minimize background and eliminate unspecific staining. The percent positivity was obtained and from four random sections within each tissue. Two-way ANOVA, followed by multiple comparisons test was performed using GraphPad Prism version 9.0, GraphPad Software, San Diego, California USA.

### Flow Cytometry

Tumors were dissociated using the MACs mouse tumor dissociation kit(Miltenyl Biotec) and suspended in PBS+1% FBS. The cells were incubated in blocking buffer containing 1:200 CD16/32(clone 93, eBioscience) for 30 min on ice. They were subsequently stained by fluorescent-conjugated primary antibodies at previously validated concentrations for 1 hr on ice. Following three PBS+1 % FBS washes, the cells were resuspended in ice cold PBS or fixed in 1 % PFA for immediate acquisition on the LSR Fortessa at the Baylor College of Medicine FACS and Cell Sorting core. Data was further analyzed using FlowJo Software version 10.0.

The following antibodies against mouse antigens were used: anti-CD3ε (145-2C11), anti-CD4 (GK1.5), anti-CD8 (53-6.7) (all from eBioscience); anti-B220 (RA3-6B2), anti-CD11b (M1/70)), anti-CD45 (30-F11), anti-F4/80 (BM8), anti-Ly6G(IA8)(all from Tonbo); anti-CD127 (SB/199, BD Biosciences), anti-CD44 (IM7, BD Biosciences), anti-CD62L (MEL14, Biolegend), Anti-KI67(16A8, Biolegend), anti-CSF1R(AFS98, Biolegend), anti-KLRG1(2F1, Biolegend).

### IMC Staining and Quantification

IMC staining and data processing was done at the Houston Methodist Immuno-Monitoring Core. Metal conjugated antibodies were ordered from Fluidigm or validated and conjugated to metals using the MaxPar antibody conjugation kit (Fluidigm). The antibodies used were suspended in BSA and azide free buffers and after conjugation they were diluted in Candor PBS Antibody Stabilization solution (Candor Bioscience) for longterm storage at 4°C.The antibodies used and metal conjugates are listed in Supplementary Excel File 2.

Paraffin embedded tumor tissue samples were sectioned onto slides with each section having a thickness of 5 um. The sections were baked at 60°C overnight, deparaffinization was performed in xylene and the sections were rehydrated in a graded series of ethanol solutions (90%, 80%, 70% and 50% for 10 min each. Antigen retrieval was performed using 1 μM sodium citrate buffer (pH 6) for 20 min in a heated water bath. Blocking buffer containing 3% bovine serum albumin in tris-buffered saline (TBS) was added to each section and the slides were incubated at R.T for 1 hr. The sections then were incubated in metal conjugated primary antibodies at 4°C overnight. The next day, samples were washed 4 times in TBS-T following which they were stained with Cell-ID Intercalator (Fluidigm) for nuclear staining. Three random sections per tumor section measuring 0.5 X 0.5 mm were ablated and imaging mass cytometry data were segmented by ilastik and CellProfiler. Histology topography cytometry analysis toolbox (HistoCAT) and R (Version 1.2.5042) were used to analyze the images and perform clustering and neighborhood analysis.

### qPCR Protocol and primer sequences

RNA was isolated from primary mouse tumor tissue using TRIzol reagent (# 15596018, Thermo Scientific). Total RNA was then reversed transcribed into cDNA using the RT2 first-strand kit (#330401, Qiagen). Realtime quantitative RT-PCR was carried out in triplicates utilizing the amfiSure qGreen Q-PCR Master Mix (#Q5602005, GenDEPOT) using a StepOnePlus real-time PCR system (Applied Biosystems, Foster City, CA, USA) and analyzed using the comparative Ct method (2-ΔΔCt) with GAPDH as the housekeeping reference gene. The primers utilized are listed in the table below.

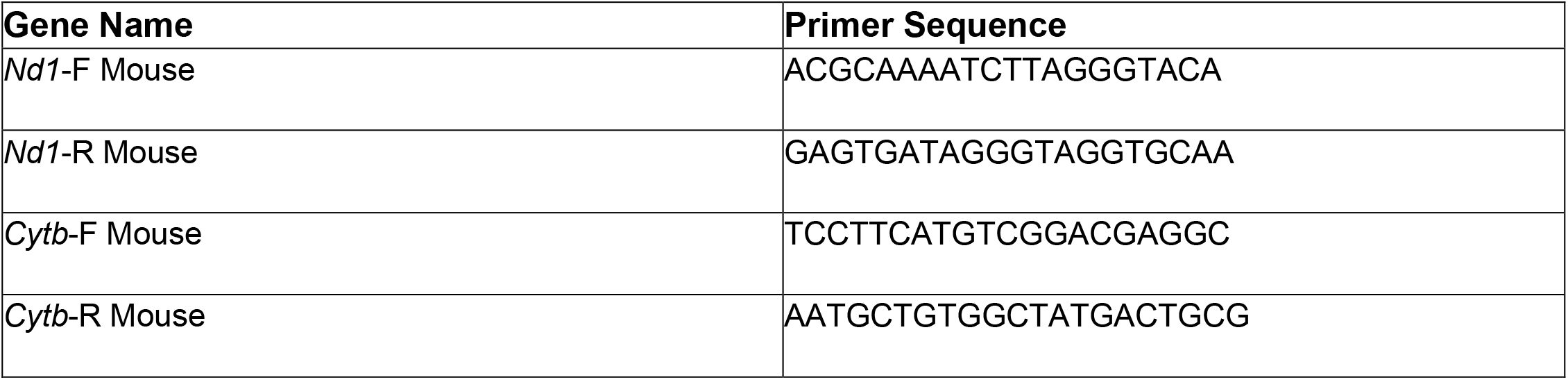

### RNA sequencing analysis

Tumor sample mRNA quality was measured using the Agilent Bioanalyzer and libraries for mRNA-seq were made using total RNA and the Illumina TruSeq mRNA sample preparation kit. Paired end (2×50bp) sequencing was performed on a Illumina HiSeq 2000/2500 sequencer at the UNC High Throughput Sequencing Facility. Consequential fastq files were aligned to the mouse mm10 reference genome using the STAR aligner algorithm. Ensuing BAM files were sorted and indexed using Samtools and quality control was performed using Picard. Transcript read counts were determined was performed using Salmon. The RNA-seq data was processed and normalized as published (2). Gene expression signatures were calculated as the median expression of all the genes in the signature as published. Non treated tumors from existing published gene expression data were used for this study (13) (and can be accessed on the Gene Expression Omnibus (GEO) under accession GSE124821. All the bulk RNA sequenc ing data for T12, T11 and 2151R is available under accession GSE173260.

### Single cell isolation and scRNA sequencing analysis

Tumors were dissociated as described previously and CD45+ cells were isolated from tumors using the Miltenyi Magnetic Enrichment Kit or FACS sorting. Fresh cells were suspended in PBS and library preparation was carried out by the Single Cell RNA Sequencing core at Baylor College of Medicine using the 10X Genomics Chromium RNA seq 5 prime Kit. The samples were sequenced at the Genomic and RNA Profiling Core at Baylor College of Medicine on Nextseq 500 or NovaSeq 6000 sequencing machines from Illumina.

Raw data has been uploaded on GEO under accession number GSE165987.

### Single cell data sequencing and processing Cellranger Count and vdj

Fastq files with range of 26-28bps for R1 and 89-96bps for R2 were processed using the Cell Ranger Single Cell Software Suite provided by 10x Genomics, V3.0.2. Reads were aligned to mouse genome (mm10) which is available on cellranger’s website http://cf.10xgenomics.com/supp/cell-exp/refdata-cellranger-mm10-3.0.0.tar.gz. Filtered gene-barcodes matrices that passes the default cellranger threshold for detection were used for further analysis. We obtained an average of 1803 unique genes per cell with a median of 1413 and an average of 7668 unique transcripts per cell with a median of 4147 which is comparable to similar scRNASeq studies. For the VDJ samples, the fastq files were processed using the cellranger vdj pipeline and the reads were alligned to the GRCm38 reference genome which is also available on cellranger’s website at http://cf.10xgenomics.com/supp/cell-vdj/refdata-cellranger-vdj-GRCm38-altsensembl-2.2.0.tar.gz.

### Data Processing

The first analysis which studies T cells consisted of four samples: 16805(PBS), 17746(CSF1R), JR20876(Combination) and JR20877(CTX). The second analysis studying Myeloid cells consisted of four samples: T12(16800 and 16805) and T11(Gao1 and Gao2) mice models. All downstream analysis were performed using Seurat R package, V3.1.3 and Monocle3_0.2.0.

### Seurat

Downstream analysis of the RNA data was performed using the Seurat R package, V3.1.3 (14). Cells with greater than 10% mitochondrial counts were removed. Outlier cells with greater than 40,000 were removed. All rps, rpl, mt and gm genes were also removed. The data was normalized and the top 3000 genes with the highest residual variance were selected as the highly variable genes seurat’s SCTransform. SCTransform also filters out genes that are present in less than 5 cells. This resulted in 31167 cells and 15976 genes from 36198 cells and 31053 genes for the T Cell study whereas it was 19352 cells and 15365 genes from 21325 cells and 31053 genes for the Myeloid study. The PCA scores were computed using Seurat’s RunPCA function and the clusters were identified using first 30PCs with a resolution of 1.2 for the T Cell study while the Myeloid study uses the same number of PCs but with a resolution of 0.8. The UMAP projection were for both studies were generated using the first 30PCs with a default parameter of 0.3 minimum distance and k = 30 neighbors.

T Cells study samples were further split into CD4 T Cells and CD8 T Cells. CD4 T Cells were subset based on Cd4 > 0.5 and Cd3e >1 while CD8 T Cells were subset based on Cd8a > 0.5 and Cd3e > 1. Cells that intersect between the CD4 and CD8 T Cells subset were removed. This resulted in a subset of 3859 unique CD4 T cells and 2346 unique CD8 T Cells and the raw counts with the original number of genes of 23393(rps, rpl, mt and gm genes were removed). SCTransform was performed on both the CD4 and CD8 T Cell subset which resulted in 3859 cells and 11758 genes while it was 2346 cells and 11918 genes for CD8 T Cell. For CD4 T Cells, the PCA scores were recalculated and the clusters were reidentified using the first 30PCs with a resolution of 1.5. For CD8 T Cells, the the clusters were identified using the first 50 recalculated PCs resolution of 1.0. The UMAP projection were regenerated using the first 30PCs for CD4 T Cells while CD8 T Cells used the first 50PCs. The other UMAP parameters remained unchanged with a default parameter of 0.3 minimum distance and k = 30 neighbors.

CD19 B cells were subset based on Cd19 > 0.5 on the SCT assay of the T Cell samples which resulted in 630 cells. SCTransform was performed on the raw counts of 23393 genes. This output a Seurat object with 630 cells and 10509 genes. PCA scores were again recalculated and the clusters were reidentified using the first 30 PCs with the default resolution of 0.8. The UMAP projections were recomputed with default parameter of 0.3 minimum distance and k = 30 neighbors.

Itgax D cells were selected by subset Itgax > 0.5 from the SCT assay of the T Cell samples. The output of the subset is 1510 cells. SCTransform was performed on the raw count of 23393 genes and this resulted in 1510 cells and 12143 genes. PCA scores were recomputed and clusters were reidentified using the first 30PCs with a default resolution of 0.8. The UMAP projections were reinitialized with the default parameter of 0.3 minimum distance and k = 30 neighbors.

For the Myeloid study, Macrophages and Monocytes cells were subset based on the SingleR Immgen annotation (15) that is explained below. 15718 cells were subsetted with the raw counts of the original 23393 genes. The data is renormalized using SCTransform which outputs 15718 cells and 14688 genes. The PCA scores were recomputed and 30PCs were used to reidentify the clusters with a resolution of 0.8. The UMAP projection were obtained using the first 30PCs with the default parameter of 0.3 minimum distance and k = 30 neighbors.

### Annotation

For CD8 T Cell, the clusters were manually annotated with the following annotation: Nfkbia+ CD8+ Tcm (Cluster 0, 6, 7, 9, 11 and 13), Ly6a CD8+ Tcm (Cluster 2), Smad7+ CD8+ Tn (Cluster 1 and 14), Gzmb+ CD8+ Teff (Cluster 4 and 5), Mki67+ CD8+ Tprof (Cluster 12) and Lag3+ CD8+ Tex (Cluster 3 and 10). As for CD4 T Cell, the clusters were manually annotated as: CD4 T naïve (Cluster 0,3,4,7,9 and 10), CD4 T cm (Cluster 1,5,8,13 and 14), Foxp3+ Tregs (Cluster 2 and 12) and CD4 T act (Cluster 6 and 11). For the CD19 B Cells, Cluster 4 and 5 were removed while the other clusters were annotated as follows:Cluster 0 and 2 - FO B Cells, Cluster 1 – Activate GC B Cells and Cluster 3 - Plasma Cells Clusters of Myeloid study and D Cells were annotated using the new SingleR, V1.0.5 which is available at https://bioconductor.org/packages/release/bioc/html/SingleR.html. The Immgen reference which has a collection of 830 microarray samples - with 20 main cell types and 253 subtypes – was used as the reference to annotate the dataset. The SingleR annotation were used to subset Myeloid cells in the Myeloid study. After the subset, the clusters were reinitialized and the UMAP were recomputed. This resulted in 20 clusters. Cluster 14 and 19 were removed due to being outliers and the rest of the clusters were annotated as: CD83+ TAMs (Cluster 1,7,10,12,13,15,17 and 18), S100a4+ TAMs (Cluster 2 and 5), Ly6c2+ Monocytes (Cluster 3 and 9), Acp5+ TAMs (Cluster 4 and 11) and Irf71+ TAMs (Cluster 0,6,8, and 16).

### Differentially Expressed Genes

Differential expression test was done using the Wilcoxon Rank Sum test which is the default of Seurat’s FindMarkers function for all genes that has not been filtered out. Log fc threshold = 0 and min.pct = 0 parameters were used. The test was conducted for both the Seurat clusters and the cell-type clusters annotated by SingleR. Three extra comparison were made for CD4 and CD8 T Cells which were Combination vs CSF1R, Combination vs CTX and Combination vs all.

### Ligand Receptor Analysis

Significant ligand-receptor (LR) pairs were identified using the same method as previously described (16, 17). We focused our analysis on the 2422 LR pairs published in these studies as well. A ligand or receptor is considered expressed if the SCT-normalized data is 0.5 in at least 10% of the cells. If a ligand or receptor is not considered expressed, the expression value will be set to zero. P-values were generated from 1,000 permutations with random shuffling the cell labels. A LR is considered significant if the interaction score is at least 2 and p < 0.01.

### Patient Data Analysis

Patient data analysis was done as described previously(28, 29). RNA sequencing and patient outcomes are available for the CALGB 40603 data in the database of Genotypes and Phenotypes (dbGaP, https://www.ncbi.nlm.nih.gov/projects/gap/cgi-bin/study.cgi?study_id=phs001863.v1.p1), while data for SCANB is available on the Gene Expression Omnibus(Accession Nos. GSE81538 and GSE96058).

### Pathway Analysis

Pathway analysis was carried out on subsets of CD68+ myeloid Cells in T12 and T11 tumors. Differential Gene Expression was obtained using the Wilcoxon Rank Sum Test, logFc > 0.5 using R studio. Signatures obtained were labeled T12vsT11 and T11vsT12 signatures (Excel, Table 3) and they were used for pathway analysis and patient data analysis in the paper. GSEA analysis (18) was used to determine if pathways were statistically different in the tumor models using FDR < 0.1 as cut off. Gene signature pathways related to metabolism were downloaded from MSigDB (19). GO and reactome analysis was performed analysis was performed using murine specific biological pathways lists downloaded from the Gene Ontology Database and Fisher’s exact test was used to determine the significance of the pathways.

### Statistical methods

Sample sizes were not predetermined for treatment studies and are denoted in the figure legends. Each in-vivo experiment was repeated at least 3 times with independent cohorts of mice. Data was show as the mean +/− the SEM. All in-vitro experiments were repeated 3 times independently. Each condition was done in biological replicates and the combined data was used to determine the P values. Statistical analysis was carried out using Graphpad Prism or R Studio (Version 1.2.5042). Statistical significance for tumor volumes was carried out using unpaired paired t-tests or two-way ANOVA (Analysis of variance) with a p value lower than 0.05 being considered significant. Survival analyses were evaluated by Kaplan-Meier curves and the log-rank (Mantel-Cox) test.

## Results

### p53 -/- syngeneic TNBC GEM models recapitulate human TNBC subtypes and respond to low dose immunostimulatory chemotherapy treatment

A previous screen using “standard of care” drugs on a claudin-low p53 null syngeneic GEM model of TNBC, T11, showed that T11 tumors were resistant to most conventional chemotherapy treatments, but responsive to low-dose cyclophosphamide (CTX,100mg/kg) (20) treatment **(Fig S1a).** We extended these studies to two other independently derived claudin-low models, T12 and 2151R. These tumors have an enrichment for the EMT pathway **(Fig 1a**) and also are highly infiltrated by TAMs **(Fig 1b, Fig S1b)** when compared to other more basal-like or luminal-like p53 -/- models such as 2225L and 2208L. Thus, they represent ideal preclinical models to develop regimens to treat the subset of TNBC that are less likely to undergo pCR following neoadjuvant chemotherapy (7). Principal component analysis showed that the three claudin-low tumor models, T11, T12 and 2151R clustered together as compared to tumor models from basal (2225L) and luminal (2208L) subtypes, indicating a certain degree of transcriptomic similarity between these models. However, there is still some level of transcriptomic heterogeneity between these three models **(Fig 1c)**. This observation led us to question whether the heterogeneity was related to the immune microenvironment in these tumors, especially in T12.

**Figure 1.**
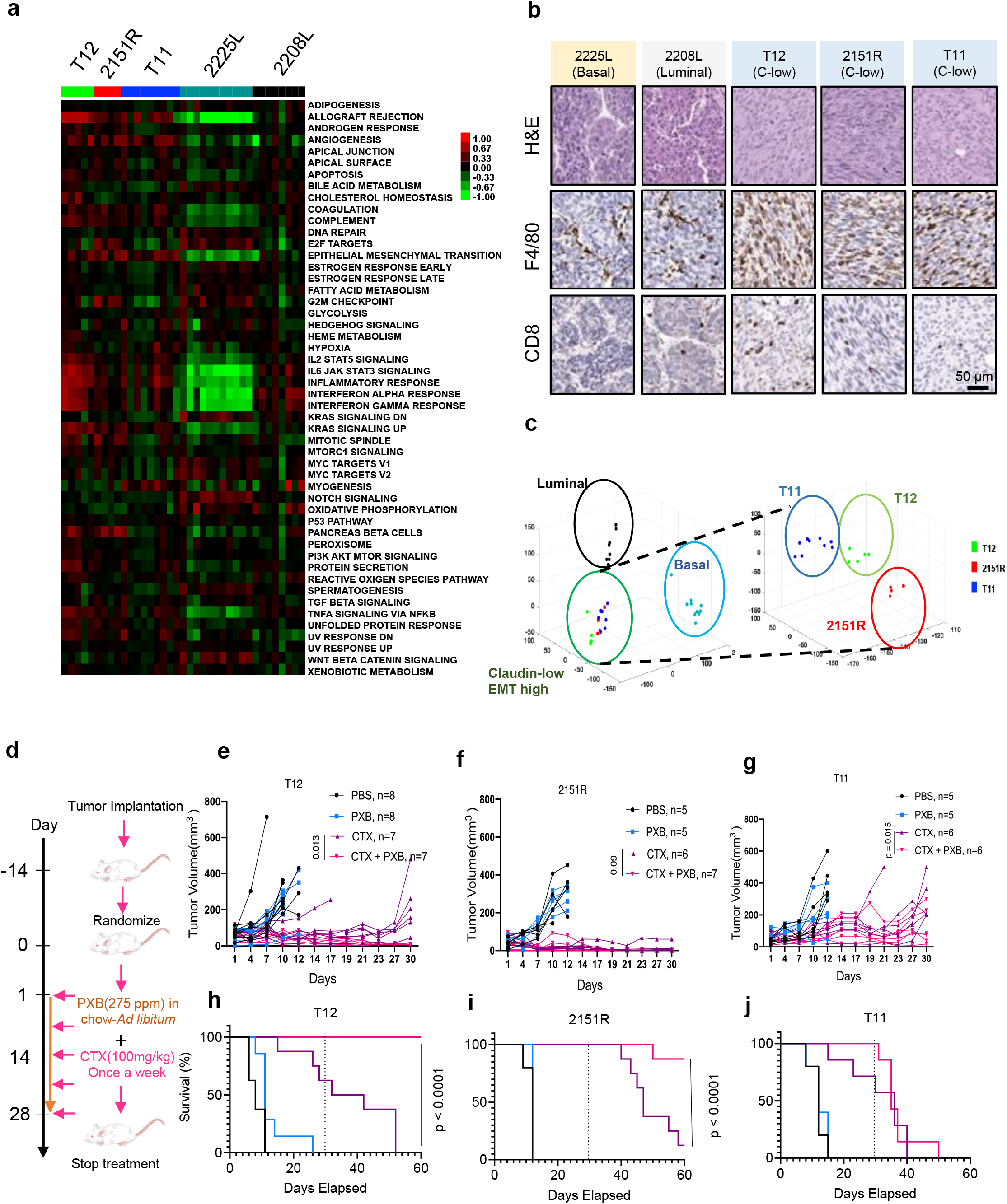
p53 -/- syngeneic TNBC GEM recapitulate human TNBC subtypes and respond to low dose immunostimulatory chemotherapy treatment. **a,** Heatmap for pathways enriched in p53-/- tumor models. **b,** Immunohistochemistry for F4/80+ TAMs and CD8+ T Cells on p53-/- tumor models. **c,** Principal component analysis (PCA) on p53 -/- tumor models. **d,** Treatment schema for combination therapy using low-dose Cyclophosphamide (CTX) and a CSF1R inhibitor Pexidartinib (PXB). **e,f,g**, Combining low-dose CTX with a Csf1r inhibitor (PXB) lead to a reduction in tumor burden in T12 and 2151R p53-/- mouse tumors and stasis in T11. Number in parentheses show the specific n values of biologically independent mice per treatment group. P value was computed by two-sided t-test. **h,i,j**, Improved survival in mouse tumor models after combination treatment. Treatment was stopped at day 30. Survival benefit was assessed by Kaplan-Meier curves using the log-rank (Mantel-Cox) test. Dotted line marks point of treatment cessation.

T12 tumors implanted in the mammary fat pads of Balb/c mice showed a greater decrease in tumor volume as compared to NSG mice that lack T Cells, B Cells and functional NK cells **(Fig S1c)** suggesting a possible synergy between T cells and CTX was required for superior T12 tumor cell killing. This was supported by in-vitro assays using GFP targeting T Cells derived from JEDI mice that showed a significant increase in cell death of GFP+ T12 cells upon addition of an active metabolite of CTX, phosphoramide mustard (PM), as CTX needs to be metabolized in the liver into its active form and cannot be used in-vitro **(Fig S1d**). Furthermore, flow cytometry analysis revealed an increase in the numbers of CD4+ and CD8+ T cells in T12 tumors after CTX treatment in immunocompetent Balb/c mice **(Fig S1e).** Antibody depletion of CD4+ resulted in an impaired response to CTX **(Fig S1f)** but did not fully abolish the differences in response between of Balb/c mice compared to NSG mice indicating there could be a possible synergy between the T cell subtypes as well as other cells NSG mice lack including B cells, that promotes tumor regression. Further analysis of treated tumors that had been treated with CD4 depleting antibody (CD4 DEP) showed an enrichment in TAMs as well as lower numbers of CD8+ T cells **(Fig S1g).** As CTX treated tumors always recurred after treatment cessation, we then asked if TAMs mediated the resistance to CTX treatment, and if so whether a treatment targeting TAMs would synergize with low-dose CTX and lead to a durable tumor response.

The enrichment of TAMs in these models **(Fig 1c)** and the previously known roles of TAM in therapeutic resistance including immunosuppression of T cell functionality (12) prompted us to hypothesize that combining CTX and anti-TAM treatment may further improve the treatment of this subset of TNBC. To test the response of these three independent models to CTX as a single agent and in combination with PXB, a small molecule inhibitor of CSF1R, we implanted the three independent claudin-low p53 null tumor models T11, T12 and 2151R **(Fig 1a, 1b, 1c)** into the mammary fat pads of immunocompetent Balb/c mice. CSF1R is a well-known macrophage recruitment and differentiation factor. Besides CSF1R, PXB also inhibits C-KIT and FLT3. It has been widely used in preclinical studies to deplete TAMs(21). Tumors were allowed to reach a size of approximately 5 mm in diameter and were then randomized into four treatment groups-PBS, CTX, PXB and CTX + PXB **(Fig 1d)**. Consistent with previous reports on other TNBC models (15), PXB failed to show efficacy as a single agent in all three claudin-low models. However, PXB had not been used in combination with CTX, which has immunostimulatory properties. All three models responded to single agent CTX treatment but recurred with differential kinetics either during treatment or following cessation of treatment. Strikingly, combination treatment led to long term durable regression in two out of three tumor models namely T12 and 2151R as well as tumor stasis in T11**(Fig 1e, 1f, 1g).** Importantly, no tumors recurred in T12 while one out of eight tumors recurred in 2151R within 30 days after treatment cessation **(Fig 1h,1i)**. In contrast, all T11 tumors grew back upon treatment cessation **(Fig 1j**). Flow cytometry analysis also revealed that combination-treated T12 tumors had a significant reduction in the numbers of F480+ CSF1R+ TAMs as compared to CTX alone**(Fig S1h)** as well as a significant enrichment of CD8+ T cells that expressed markers of central memory T cells (CD62l and CD127) and CD8+ effector memory T cells (CD62L+CD127+CD44+), as compared to single agent-treated tumors (**Fig S1i).** This therapeutic combination was also successful with a 3.3X lower dose of PXB **(Fig S1j).** Finally, we also used a high affinity monoclonal antibody, Axatilimab (22), directed towards CSF1R to treat T12 and 2151R tumors in combination with low-dose CTX. Combination of Axatilimab and CTX lead to dramatic tumor regression in both models and depleted F480+ macrophages within the tumor in T12 tumors (**Fig S1k, S1l**).

### Combination therapy leads to an expansion of CD8+/CD4+ T cells and B cells in responsive T12 tumors

We next employed imaging mass spectrometry to visualize and quantify the spatial interactions of various immune cell types in the tumor immune microenvironment in both the T12 highly responsive and T11 poorly responsive models (**Fig. 2a and 2b**). Compared to treatment of PBS or single agents PXB or CTX, the CTX + PXB combination increased juxtaposition between T cells (**Clusters 1-7**) and B cells (**Cluster 14**) or dendritic cells (**Cluster 15**). Interestingly, CTX alone appeared to stimulate interactions between T cells and macrophages (**Cluster 18**). Addition of PXB diminished this effect. The increased interactions between T cells and macrophages may potentially explain the reduced response to single agent CTX.

**Figure 2.**
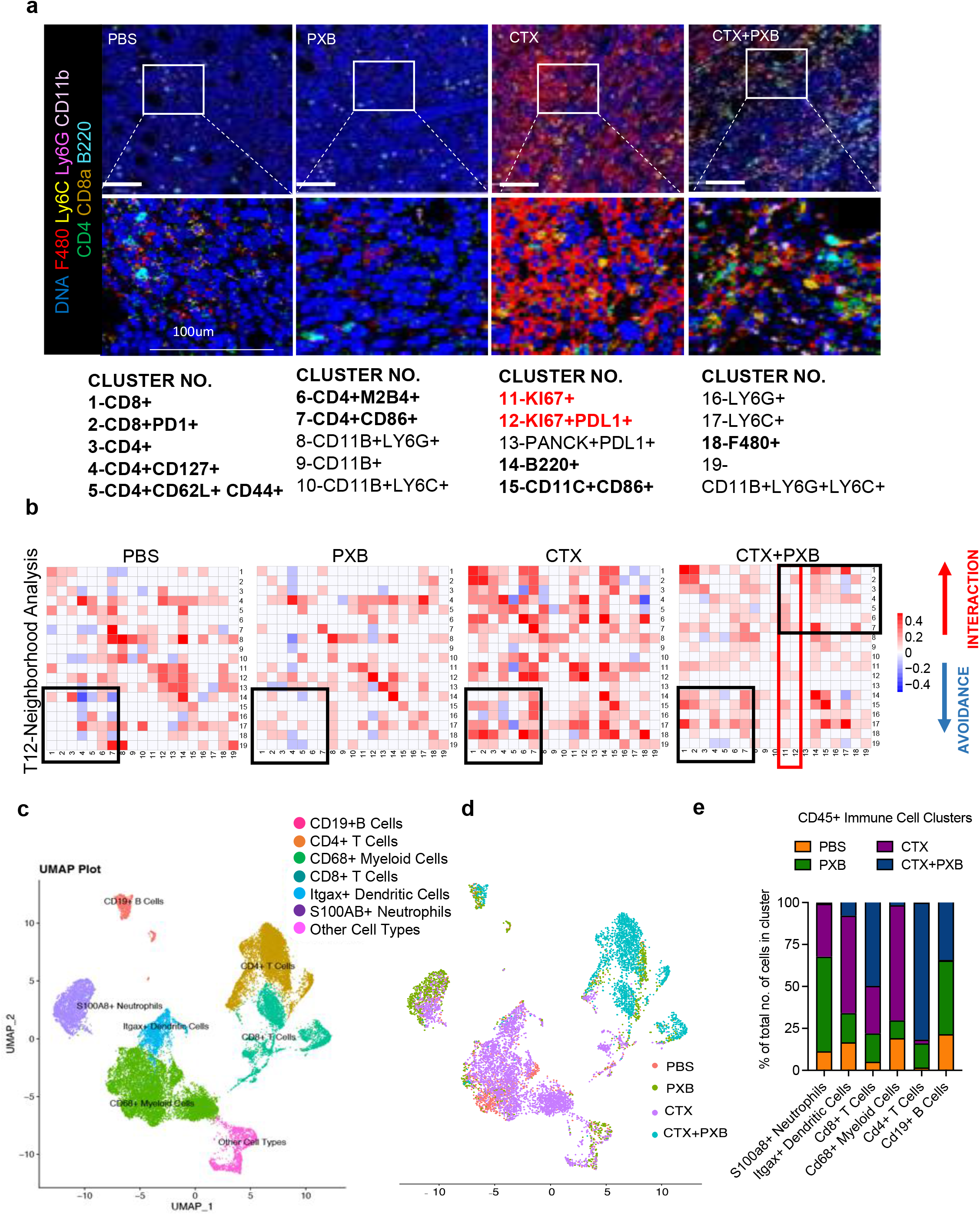
Combination therapy leads to an expansion of CD8+/CD4+ T cells and B cells in responsive T12 tumors. **a**, Imaging mass cytometry analysis of the tumor immune microenvironment in T12 tumors before and after treatment. Representative images overlaid with 7 markers (F480, Ly6C, Ly6G, CD11b, CD4, CD8a, B220) for each treatment group. **b,** Neighborhood analysis of T12 tumors in which the color of the squares indicates significant pairwise interactions or avoidance between PhenoGraph defined cellular metaclusters. Highlighted interactions include CD8+/CD4+ T cells (clusters 1-7), B220+ B cells (cluster 14) and CD11C+ CD86+ Dendritic cells (cluster 15). Three Regions Of Interest (ROI) were ablated per tumor section. n= 3-5 independent biological replicates per treatment group. **c,** UMAP plot showing CD45+ immune cell clusters before and after treatment in T12 tumor model. **d,** UMAP plot showing CD45+ immune cells split by treatment group. **e,** Quantification of main immune cell clusters in 4 treatment groups.

T11 tumors that responded poorly to combination therapy were populated by Pan-CK+ KI67+ PDL1 + proliferating **(Clusters 11 and 12, Fig S2a, S2b**) tumor cells that had no significant interactions with with subpopulations of CD8+ and CD4+ T cells **(Clusters 1-7, Fig S2a, S2b**) indicating the lack of infiltrating T cells unto the tumor core. T cell and B cell interactions were also not observed in T11 tumors after combination therapy, contrasting with T12. This is indicated by **Clusters 1-7 and 14 (Fig S2a, S2b-highlighted in black**). As indicated by clusters 11 and 12 **(Fig 2b-highlighted in red**), there was a reduced number of Ki67+ proliferating cells that were in close contact with CD4+ T cells in T12 tumors following combination therapy **(Cluster 4,5,6 and 7, Fig 2b-highlighted in black)**. Studies have shown that TAMs can be recruited by tumors cells to escape T cell mediated killing(12) using Flow cytometry, we found that the number of CSF1R+ F480+ TAMs after combination treatment was higher in non-responsive T11 tumors **(Fig S2d).** TAMs have immunosuppressive effects on T cell mediated tumor cell killing, T cell differentiation and proliferation, which may explain the resistance of T11 tumors to combination treatment.

To further elucidate the immune effects of CTX + PXB in the highly responsive T12 TNBC model, we performed scRNA-seq and combined V(D)J sequencing of T cell receptors (TCR) on CD45+ immune cells across all four treatment groups. Uniform Manifold Approximation and Projection (UMAP) projections of the cells revealed different clusters of immune cells across the four different treatment groups including *Cd68+* TAMs, *S100a8+* Neutrophils, *Cd19+* B cells as well as *Cd8+* and *Cd4+* T cells **(Fig 2c, S2c, S2e**). However, most immune cells from PBS and CTX treated mice clustered separately from combination-treated tumors **(Fig 2d).** Upon further analysis, we observed a nearly complete depletion of *Cd68+* TAMs and low *S100a8+* neutrophils in the combination-treated group as well as a significant expansion of *Cd4+* and *Cd8+* T cells. The expression of common TAM marker gene *Adgre1(F480),* was not expressed at a consistent/detectable level on shallow scRNA sequencing and so *<u>Cd68</u>* was used as a monocyte/TAM marker. PXB single agent-treated tumors had high levels of *S100a8+* neutrophils supporting previous studies showing the accumulation of tumor infiltrating neutrophils (TINs) upon TAM depletion in treatment refractory mice **(Fig 2d, S2f)** (12). Pathway analysis using genes significantly enriched in the *S100a8+* neutrophil cluster as compared to the other immune cells, showed a significant enrichment of pathways related to granulocyte and monocyte activation and aggregation, and pathways related to the suppression of T cell proliferation and activation **(Fig S2g**).

### TAMs in responsive T12 and non-responsive T11 tumors are phenotypically distinct

The lack of durable response in T11 tumors was unexpected and led to the question why these genetically and phenotypically similar models displayed a differential response to combination treatment. To address this question, we further analyzed the scRNA-seq data using Seurat. Five major clusters of TAMs were identified, including *Ly6c2+* Monocytes, *Cd83+* TAMs, *Irf7+* TAMs, *Acp5+* TAMs and *S100a4+* TAMs **(Fig 3a, 3b, S3a**). Strikingly, T12 TAMs are predominantly *Cd83+* whereas T11 TAMs appear to be more heterogeneous and distribute in all five clusters. Interestingly, the baseline expression of *Csf1r* was significantly higher in T12 than T11 TAMs, suggesting a lower dependence on this signaling pathway by T11 TAMs for survival/differentiation (**Fig 3c, 3d).**

**Figure 3.**
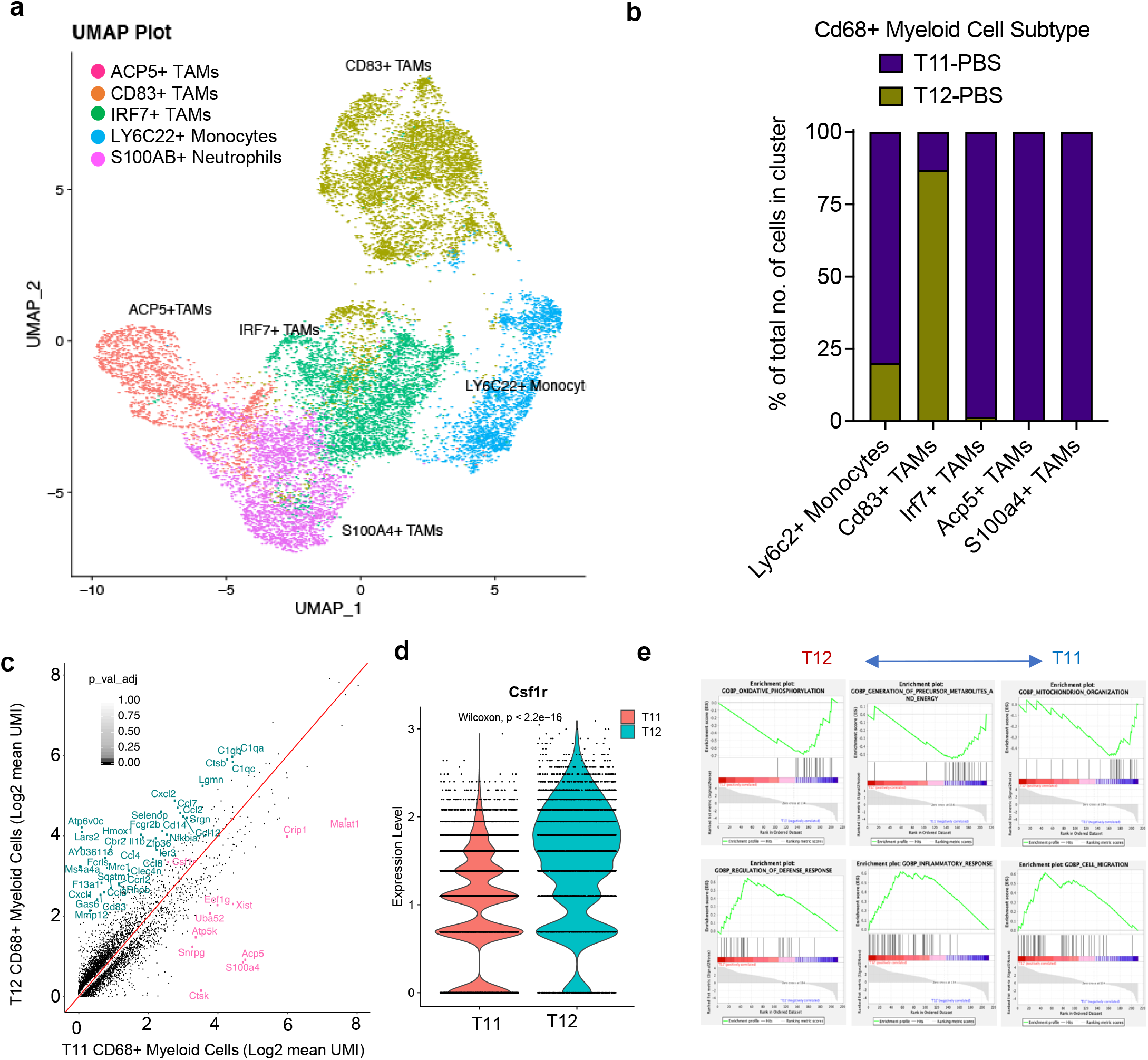
TAMs in responsive (T12) and non-responsive (T11) tumors are phenotypically distinct. **a,** UMAP plot showing CD68+ Myeloid cells sub-clysters including *Ly6c2+* Monocytes, *Irf7+* TAMS, *Acp5+* TAMS and *S100a4+* TAMS from untreated T12 and T11 tumors (2 per group, Gene names are italicized and protein names are in uppercase). **b,** Quantification of TAM sub-clusters in untreated T12 and T11 tumors. **c,** Plot showing differentially expressed genes between T12 TAMs and T11 TAMs (log2fold change>0.5, p < 0.05). **d,** Quantification of Csf1r expression in T12 and T11 TAMS. P values > 0.05 were considered significant and calculated using Wilcoxan Rank Sum Test. **e,** Enrichment of select activation and metabolism related pathways in T12 and T11 CD68+ Myeloid cells using GSEA.

In fact, we identified a G12V activating mutation of k*ras* in T11 (**data not shown**), which was accompanied by increased pMAPK compared to T12 **(Fig S2h),** all of which may contribute to more resistance to combination treatment of T11 tumors.

To further delineate the differences between T11 and T12 TAMs, we performed pathway analysis on differentially expressed genes **(logFc> 0.5, Fig 3e**) in pre-treatment T11/T12 TAMs. T12 TAMs were enriched in pathways related to immune activation including those involved in defense and inflammatory responses and TAM activation markers such as C1qa **(Fig S3b)**. They also showed a downregulation of genes related to oxidative phosphorylation(ox-phos**) (Fig S3c).** In contrast T11 TAMs expressed higher levels of oxidative phosphorylation related genes such as *CytB* and *Nd1* **(Fig S3d).** Interestingly, 2151R and T12 had similar levels of expression of these genes at a transcript level, which suggests a degree of metabolic similarity in these two responsive models. Surprisingly, T12 TAMs expressed higher levels of markers related to immunosuppressive macrophages such as *Spp1* **(Fig S3b).** This indicates that M2 “like” T12 TAMs are also sensitive to CSF1R inhibition. Further studies are required to elucidate the role of metabolism in TAMs that survive CSF1R inhibition and whether targeting oxidative phosphorylation could provide a vulnerability to inhibit these cells.

### Combination therapy leads to polyclonal expansion T cells that exhibit memory cell phenotypes

To further understand T cell functionality in the absence of immunosuppressive tumor infiltrating TAMs after combination treatment, we next focused our analysis on *Cd8+* T cells in all four treatment groups. The reclustering of 2346 CD8+ T cells (Methods) from all four treatment groups in T12 tumors revealed six different clusters of T cells including *Smad7+* Naïve *Cd8+* T cells, *Nfκbia+ Cd8+* central memory/memory precursors T cells, *Ly6a+ Cd8+* central memory T, *Gzmb+ Cd8+* effector T cells, *Mki67+ Cd8+* proliferative T cells, and *Lag3+* exhausted *Cd8+* T cells **(Fig 4a, S4a).** The *NFκbia+Cd8+* central memory T cells (Tcm) were greatly increased after combination therapy as compared to the other sub-populations of T cells **(Fig 4b)**. These cells in the combination treated group expressed high levels of markers such as *Jun, Id3* and *Nfxbia* which are markers of exhaustion resistance(23), longevity(24) and are required for the maintenance of secondary lymphoid structures(25). Additionally, these cells expressed markers of early activation such as *Cd28* and *Cd69* that also control T cell differentiation as shown in the corresponding feature plots. Consistent with these results, combination treated *Cd8+* T cells also had a low expression of terminal exhaustion markers such as *Pdcd1, Lag3* and *Tigit* as compared to those in untreated or single agent treated mice **(Fig 4c, S4b**). Thus, scRNA-seq analyses confirmed flow cytometry results and revealed a profound expansion of long-lived, *Cd8 +* memory T cells upon combined PXB and CTX treatment, which may help explain that durable responses were observed for up to 30 days post-treatment.

**Figure 4.**
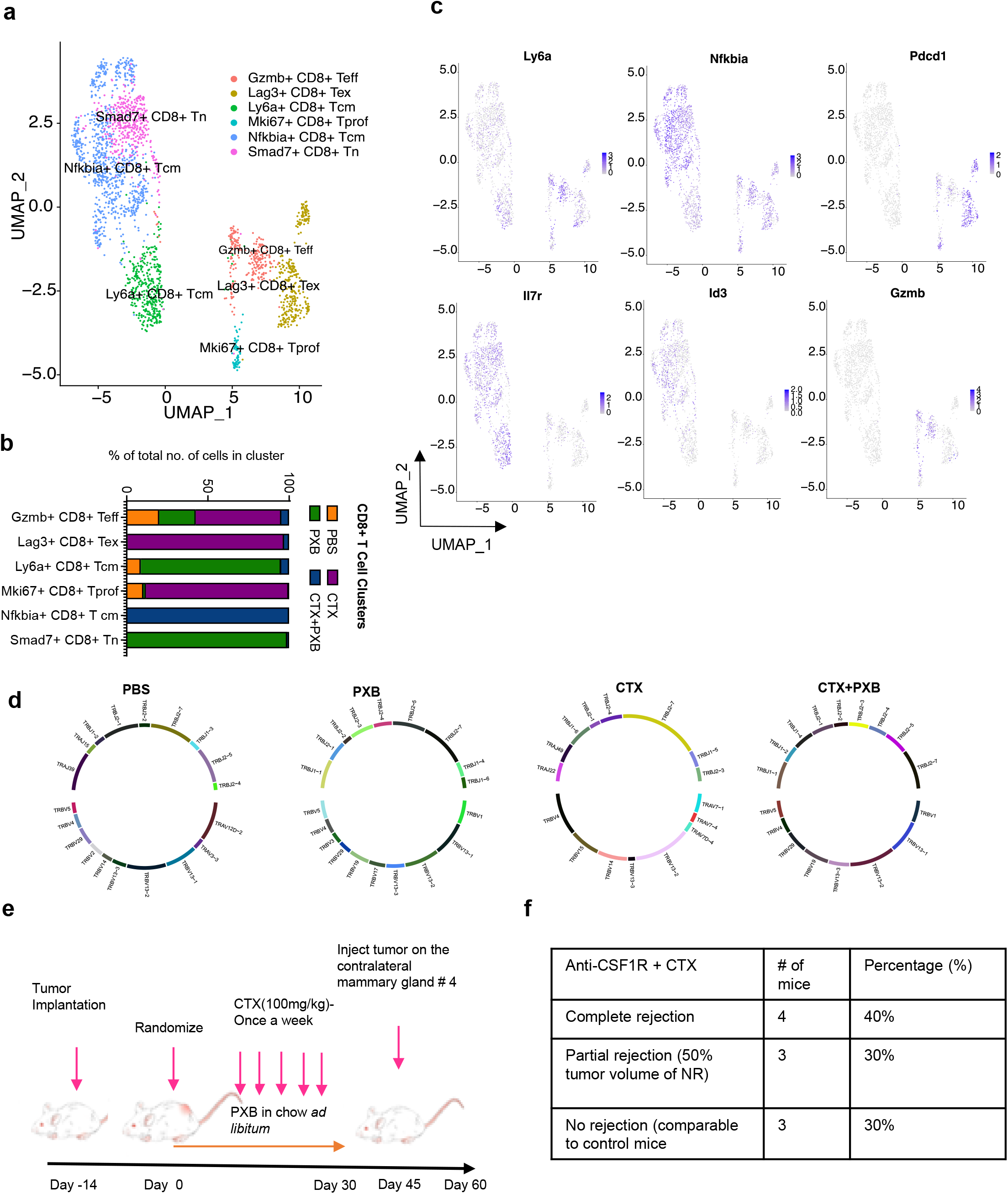
Combination therapy leads to polyclonal expansion T cells that exhibit memory cell phenotypes. **a,** UMAP plot showing *Cd8+* T Cell clusters of the 4 treatment groups and split by sample type. Clusters were annotated using the Immgen database (5), SingleR (Data not shown) and using known markers. **b,** Quantification of *Cd8+* T cell subsets in different treatment groups. **c,** Feature plots showing the expression of select memory/exhaustion/activation markers in CD8+ T cells. **d,** Clonal frequency of *Cd8+* T cells in all 4 treatment groups and chord diagrams representing unique V-J region pairings in *Cd8+* T cells in T12 tumors before and after treatment. **e,** Schema for T12 rechallenge experiments and treatment. **f,** Quantification of mice that completely or partially (tumor volume < 50 % of control tumors) rejected T12 mammary tumors injected into the contralateral mammary gland of previously treated T12 tumor bearing mice, n=10.

Most anti-tumor responses by CD8+ T cells are due to their clonal expansion triggered by specific tumor antigens that are presented by antigen presenting cells (APCs) such as CD11C+ dendritic cells and B220+ B cells(26). Macrophages can present antigens as well but cannot infiltrate lymph nodes or secondary lymphoid structures that are usually the sites of antigen presentation. In these experiments, exhausted CD8+ T cells in PBS- and CTX-treated groups exhibited higher clonality, which is consistent with studies showing monoclonal enrichment of exhausted T cells after checkpoint blockade(27). However, the T cells in the combination-treated group showed a polyclonal expansion with no enrichment for a single clone **(Fig 4d, S4c),** suggesting that depletion of TAMs may fundamentally alter the antigen-presentation and/or clonal expansion process.

To functionally validate the memory status of T cells after combination treatment, we re-challenged T12 tumor-bearing mice 15 days post-treatment with fresh T12 tumor chunks that were implanted into the contralateral mammary gland. Seventy percent of these mice showed a complete or partial rejection of these newly implanted tumors **(Fig 4e, 4f).** This result further supports the role of the combination treatment on establishing long-term immune memory against tumor cells.

Using combination treatment, we were also able to successfully target established T12 lung metastases using an experimental metastasis tail vein model (**Fig S4d).** CTX alone was also able to successfully inhibit lung metastasis as compared to PBS or PXB treated mice **(Fig S4e**). Single agent PXB treatment significantly increased the lung metastasis burden, when compared to PBS-treated mice **(Fig S4f, S4g).** These results are like those reported for MDA-MB231 cells in SCID mice indicative of a distinct lung tumor microenvironment where a subset of TAMs could play a stronger anti-tumor role as compared to the TME in the mammary gland tumors (28). CTX alone also lead to a reduction of lung metastasis however combination treatment resulted in a significantly higher number of B220+ B cells and CD4+/CD8+ T cells as compared to CTX alone **(Fig S4h, S4i).** Follow up studies are needed to test whether combination treatment can promote long term regression of lung metastasis as compared to CTX alone.

For CD4 T cells treatment, UMAP analysis showed four different clusters *Ccr7+ Cd4+* naïve, *Cd40lg+ Cd4+* Tcm, *Cd69+ Cd4+* T cells(activated) and *Foxp3+* T regulatory cells. Out of these *Cd4+* T naïve (Tn) and *Cd4+* Tcm cells were enriched after combination therapy while Foxp3+ T regs were enriched in single agent CTX treated mice **(Fig 5a, 5b)**. The *Cd4+* Tn cells expressed common naïve markers such as *Ccr7, Sell, Lef1* and low levels of *Cd69* which is typically a marker of early T cell activation(29). However, CD4+ Tcm cells enriched after combination treatment has increased expression of *Cd69* and other markers associated with T cell longevity and survival such as *Id3/Fos/Jun* **(Fig 5c, S5a, S5b)**.This supports recent studies that have shown that the transcriptomes of Tn and Tcm cells derived from patients do not represent two discrete states between these two populations but rather show a continuum of gene expression(30).

**Figure 5.**
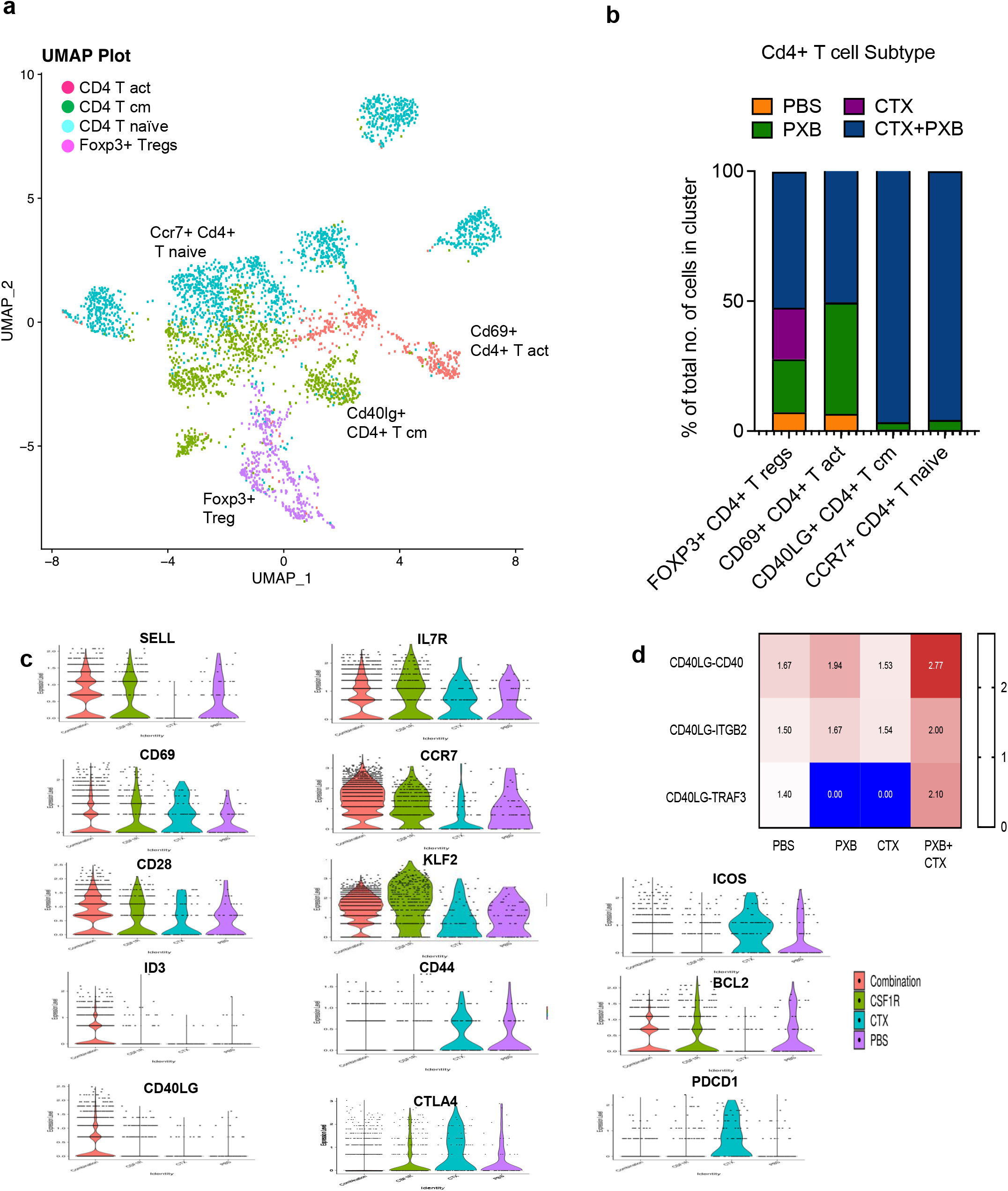
CD4+ T cell and B cell play an important role in mediating long term tumor regression. **a,** UMAP plot showing *Cd4+* T Cell clusters in the 4 treatment groups. Clusters were annotated using the Immgen database (5), SingleR (Data not shown) and using known markers. **b,** Quantification of *Cd4+* T cell subsets in different treatment groups. **c,** Violin plots showing the expression of select memory, activation and exhaustion markers in *Cd4+* T cell sub-clusters. **d,** select ligand-receptor pairs involved in *Cd4+* T cell and *Cd19+* B cell signaling. P values < 0.01 were considered significant (See Methods). List of significant ligand receptor analysis between *Cd4+* T cells and *Cd19+* B cells, *Cd68+* Myeloid Cells and *Itgax+* Dendritic cells is available in the supplementary materials.

Additionally, V(D)J analysis revealed a polyclonal expansion of these subsets as well **(Fig S5c)** *Cd4 +*T naive and T cm cells expanded after combination treatment also displayed increased expression of the CD40 ligand (lg)which is a member of the Tumor Necrosis Factor (TNF) family of ligands **(Fig 5c)**. CD40lg is expressed primarily on activated T cells. However, it is also present on B cells, dendritic cells, and macrophages. The receptor for CD40LG is CD40. It was first discovered on B cells but is also expressed on other antigen presenting cells such as dendritic cells (31). CD40/CD40lg is essential for the survival of antigen presenting cells including B cells where it can lead to the maturation of B cells, increase their ability to effectively present antigens to T cells and form germinal centers (32). To determine if there was a relationship between the *Cd40lg* Tcm cells and the *Cd19+* B cells that expanded after combination treatment we performed ligand receptor analysis for all four treatment groups. Interestingly, we identified a unique interaction between CD4 T cells and B cells in the combination treated group which was *CD40lg* and *Traf3* **(Fig 5d).** Previous studies have implicated TRAF3 in the inhibition of canonical and non-canonical NF-κB signaling as well as a reduction in CD40 transcript. However, this effect is thought to be cell type specific. Other studies have shown that TRAF3 can mediate class switching which is essential for B cells to proliferate and generate a diverse antibody repertoire directed towards multiple tumor antigens (33). Based on these results we then decided to study the reciprocal interactions between B cells and CD4 Tcm cells in the singly and combination-treated mice.

### Activated B cells expand after combination therapy and may be the main APCs within the tertiary lymphoid structures

scRNA-seq data for all four treatment groups for *Cd19+* B cells revealed three discrete subclusters including *Cd19+* B cells, *Cd86+* activated B cells (ABCs) that expressed high levels of Cd40 and Germinal center (GC)-like B cells that expressed GC markers such as *Bcl6*. ABCs were enriched after combination treatment **(Fig 6a, 6b)** and these cells had increased expression of *Cd40* and *Rel* as compared to the naïve B cells **(Fig S6a**). Rel is a member of the NF-κB family of transcription factors and is essential **f**or the long-term survival of ABCs. These B cells also expressed immunoglobulin transcript genes such as *Ighm* and *Ighd* **(Fig S6a),** however this remains to be validated at a protein level.

**Figure 6.**
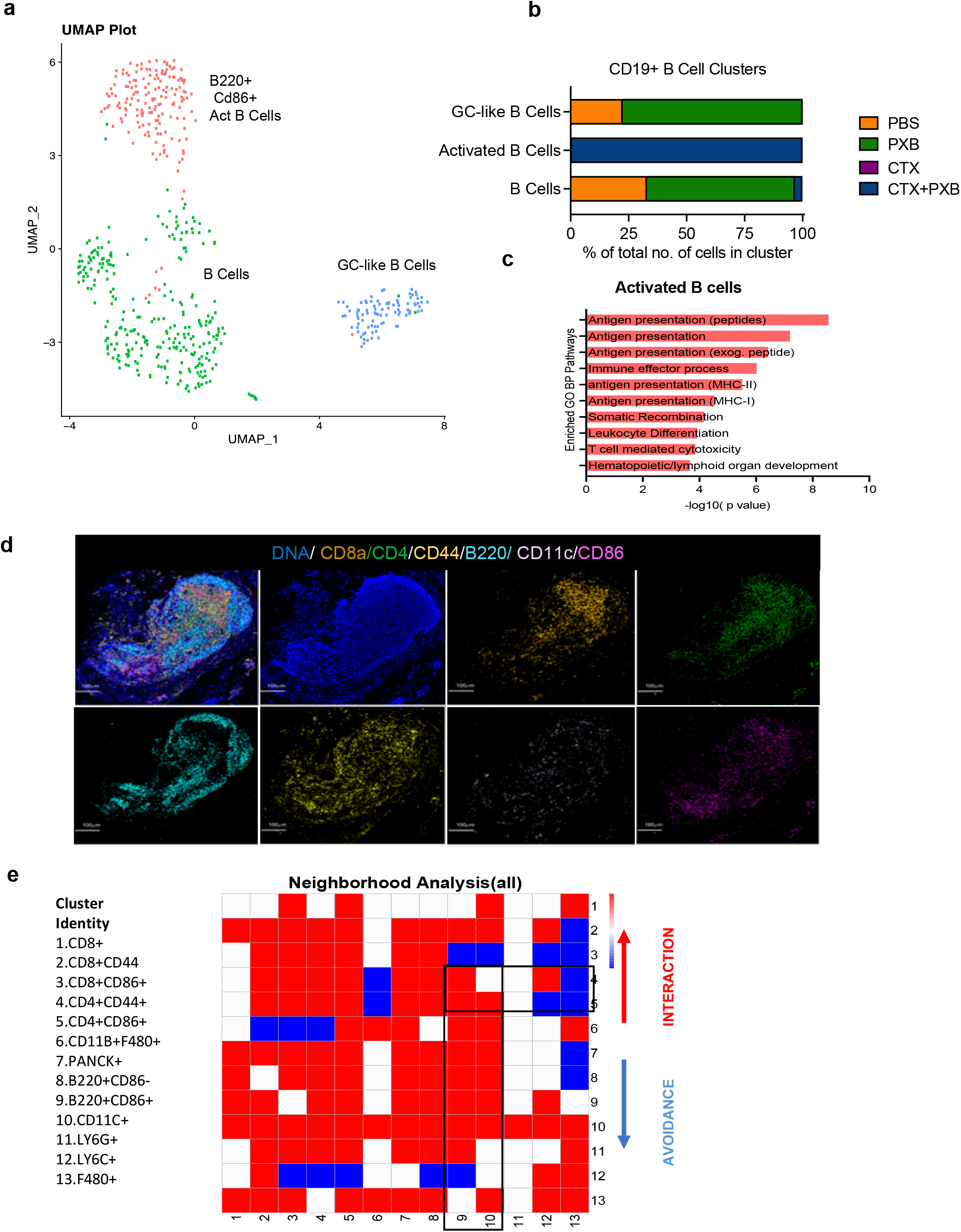
Activated B cells expand after combination immunotherapy and are the main antigen presenting cells to CD4+ T memory cells with tertiary lymphoid structures. **a,** UMAP plot showing *Cd19*+ B Cell clusters of the 4 treatment groups and split by sample type. Clusters were annotated using the Immgen database (5), SingleR (Data not shown) and using known markers. **b,** Quantification of *Cd19+* B cell subsets in different treatment groups. **c,** Pathway analysis showing select enriched biological process pathways in activated CD86+ B cells that expand after combination treatment. **d,** Imaging Mass Cytometry analysis of TLS observed 30 days post-treatment in long-term responder mice bearing T12 tumors. Representative image overlaid with 4 markers (F480, CD8A, B220, CD4, CD44, CD11C, CD86) in the site of completely regressed primary tumors in combination treated mice 30 days post-treatment. **e,** Neighborhood analysis of tertiary lymphoid structures in which the color of the squares indicates significant pairwise interactions (Red), avoidance (Blue) or a complete lack of interactions (White) between PhenoGraph defined cellular metaclusters. Highlighted interactions include CD4+/CD44+ memory T cells cells (clusters 4-5) with B220+/CD86+ B cells (cluster 9) and CD11C+ Dendritic cells (cluster 10).

Besides CSF1R, PXB has also been shown to inhibit FLT3, which is an important dendritic cell (DC) maturation factor. We did not observe an expansion of antigen presenting DCs after combination treatment. However, pathway analysis of ABCs showed increased expression of antigen presentation pathways to both CD8+ and CD4+ T cells indicating that after combination treatment, B cells most likely are the chief antigen presenting cells instead of dendritic cells or macrophages **(Fig 6c**). Reactome analysis showed an upregulation of NF-κB survival signaling indicating the *Cd40lg-Traf3* interactions are not inhibitory in this context **(Fig S6b**). These data suggest that both CD4+ T cells and CD86+ activated B cells are essential for long-term response to CTX and PXB in our models.

Consistent with this long-lasting durable response to combination treatment, we observed presence of TLSs in the tumor beds of treated mice bearing regressed T12 tumors six weeks post-treatment as compared to the untreated or single agent tumors that were harvested seven days post-treatment **(Fig S6e)**. We also observed TLSs in the primary mammary gland site in mice that completely rejected T12 tumor rechallenge **(S6c)**. Furthermore, they expressed endomucin, which is a component of heavy endothelial venules(34)-a known TLS marker **(Fig S6d).** Since long lasting TLS have been implicated in improved responses to checkpoint blockade therapy, we performed IMC and observed highly significant interactions between subsets of CD8+/CD4+ memory T cells **(Clusters 1-5, Fig 6e)** and B220+ B cells **(Clusters 8 and 9, Fig 6e)** and avoidance of CD11B+ F480+ macrophages and LY6C+ monocytes **(Clusters12 and 13, Fig 6e)**. Using markers obtained from our scRNA-seq analysis to further delineate B cell heterogeneity at a protein level, we identified two B cell subpopulations which were B220+CD86+ and B220+CD86**-(Clusters 8 and 9, Fig 6e**).

We next asked if whether a certain B cell subset preferentially interacts with T cells in TLS. We observed significant interactions between B220+ CD86+ B cells and CD8+/CD4 + CD44+ T cells **(Clusters 1,2,4,5 with cluster 9, Fig 6d)**. CD44 is a known memory T cell marker along with CD62l, but we did not detect a high number of these cells in our scRNA-seq data while the mice were on treatment and only identified these cells in TLS, 30 days post therapy. They could possibly represent a subset of T cells that expand after combination therapy and are long lived/help in promoting long-term tumor regression. CD4+CD44+ T cells **(Cluster 4)** interacted with B220+CD86+ B cells **(Cluster 9)** but had no significant interactions with CD11C+ Dendritic Cells **(Cluster 10, Fig 6e)**. This indicates the possibility that antigen presenting B cells can persist at least 6 weeks post-treatment and may be the main APCs in long-lasting TLSs after combination therapy. Interestingly, one of the top pathways identified in the GO analysis of activated B cells in combination treated mice was related to lymphoid organ development **(Fig 6c).** Further studies are needed both to establish a functional role and to understand the mechanisms by which B cells might promote the formation of anti-tumor TLSs.

### T12 TAM signature is upregulated in patients with Claudin-low breast cancer

To determine if there was any clinical correlation with these immune cell subsets, we derived a signature using differentially expressed genes (log Fc>0.03) from the T12 and T11 tams, which are included in Supplemental Table S1. We applied T12 TAM and T11 TAM signatures to two data obtained from stage II-III TNBC patients from the CALGB 40603 trial (35) and the SCAN-B dataset (36). The T12 TAM signature was significantly upregulated in human claudin-low tumors from TNBC patients, while the T11 TAM signature was not specifically associated with these human claudin-low tumors (**Fig 7a, 7b, 7c, 7d).** Additionally, high expression of the T12 TAM signature was associated with a decreased overall survival in TNBC patients from the SCAN-B dataset **(Fig 7e).** We also saw the increased expression of the T12 TAM signature when we looked at claudin-low tumors across all breast cancer subtypes including TNBC in multiple datasets including TCGA, METABRIC and SCANB **(Fig S7a, S7b).** This underscores the complexity of TAMs even within closely related claudin-low murine models and establishes the T12 murine model that show similar tumor gene expression (37), and now here, similar immune cell features as human claudin-low tumors.

**Figure 7.**
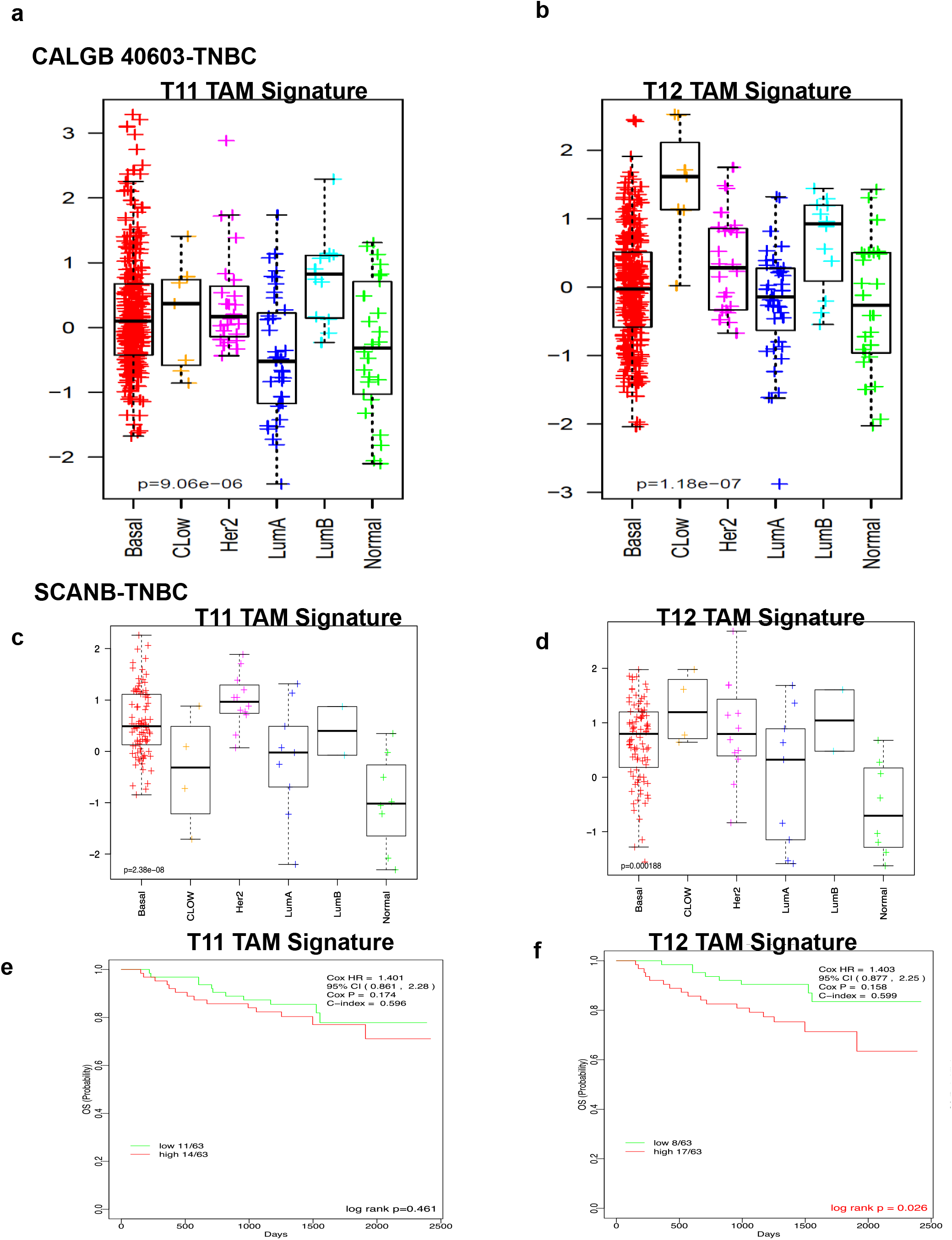
T12 TAM signature is upregulated in patients with Claudin-low breast cancer and activated B cell signatures correlate with pCR in TNBC patients after neoadjuvant therapy. **a,** Downregulation of T11 TAM signature in TNBC claudin-low breast cancer patients as compared to other TNBC subtypes in CALGB 40603 and SCANB clinical trial datasets. **b,** Upregulation of T12 TAM signature in TNBC claudin-low breast cancer patients as compared to other TNBC subtypes in CALGB 40603 and SCANB clinical trial datasets. **c,** Association of T12 TAM signature with decreased overall survival in TNBC patients from the SCANB dataset.

## Discussion

In this study, we performed an in-depth analysis of single agent and combination treated preclinical TNBC models using the latest single cell RNA and VDJ-seq techniques, as well as sophisticated imaging technologies such as IMC. These tools have provided new insights into changes in the immune microenvironment before and after combination treatment with both chemo- and macrophage-targeting therapies. Surprisingly, these studies revealed a subtle heterogeneity that exists between genetically and phenotypically related models that appear to be enriched for similar immune cells types such as macrophages and pathways such as the EMT pathway. The models used in this study, namely T12, T11 and 2151R have shown varying responses to a low dose CTX and a small molecule CSF1R inhibitor. While T12 and 2151R models display a dramatic response leading to as complete long-lasting regression of the primary tumors upon treatment cessation, T11 tumors display stasis while undergoing treatment but recurred rapidly once treatment cessation.

Spatial analysis done in TNBC has shown that tumors with a high number of CD8+ T cells that infiltrate into the tumor core have more favorable prognosis as compared to stroma restricted cells, indicating the importance of understanding the TIME architecture with respect to immune cells (16). One limitation of IMC analysis is the requirement to pick random portions of the tumor section to ablate and visualize. Thus, IMC is not the best method to quantify immune cell numbers as compared to flow cytometry or immunohistochemical staining techniques such as IF/IHC, where you can image the entire section, especially for large untreated tumor sections in the untreated groups. This is illustrated in the staining for F480+ TAMs in PBS treated tumors. However, the power of this technique enabled us to visualize the spatial interactions between a large number of tumor infiltrating cells including Ki67+ tumor cells and a variety of immune cells.

TAMs classically have been classified as either M1-like and M2-like type macrophages with activating and immunosuppressive functions, respectively although more recent studies have illustrated that this designation is overly simplified and there is a much more complex spectrum of activities. Studies that have performed scRNA sequencing on patients with colorectal cancer have identified subsets of C1QB+ macrophages that have differential sensitivity to CSF1R blockade(38). These results are consistent with our data that shows T12 TAMs have high *C1qb* expression, also express higher levels of *Csf1r* and are more sensitive to PXB. Conversely the T11 TNBC model is infiltrated by highly metabolically active TAMs that are marked by the expression of *S100a4* and *Acp5.* Additionally, while T12 TAMS appear to have an increase of the cytokine metabolism pathways and genes related to glycolysis, T11 TAMs seem to depend upon oxidative phosphorylation for their energy needs. Whether this is a consequence of the *kras* mutation observed in the T11 model remains to be established.

Depletion of TAMs within the tumor microenvironment coupled with a low dose of immunostimulatory chemotherapy was able to elicit durable tumor regression and an expansion of polyclonal long-lived central memory T cells in certain TAM infiltrated models. These cells persist for long periods of time at the primary tumor site after tumor regression and they may also facilitate tumor rejection upon rechallenge. This response was dependent upon the combination of the immunostimulatory chemotherapy with CSF1R inhibition and was not observed with single agents. Previous studies with PXB in combination with a taxane were performed using the MMTV-PyMT model which is classified as a luminal-like breast cancer model (21). These studies focused on the role of CD8 T cells and targeting macrophage recruitment/response pathways in combination with cytotoxic chemotherapy, but durable responses were not observed in this model. Furthermore, these studies did not report a role for B cells or the spatial relationships between B and T cells. The present study identified mouse models that will provide invaluable tools to study TAM heterogeneity in the future. Furthermore, the publicly available scRNA seq datasets of T11 and T12 tumors provide a resource to explore TAM heterogeneity in greater detail. It also uncovered the presence of a highly conserved TAM signature derived from the T12 claudin-low tumor model across multiple human breast cancer datasets including TNBC and provides us with a pre-clinical rationale to test out a combination of CTX and anti-CSF1R inhibition in the clinical setting. Further studies are required to identify therapeutic vulnerabilities in the poorly responsive T11 claudin-low model as well as identify CSF1R inhibitors with lower levels of toxicity and higher specificity.

Another unexpected and unique observation seen in the present studies was the existence of TLSs within the primary tumor site that persisted post treatment. While these structures have been implicated in improving the prognosis for patients that respond to checkpoint inhibition, they are usually thought to be transient and have not been studied in detail in mouse models of TNBC. We observed that TLS co-infiltrated by antigen presenting CD86+ B cells and CD4+ CD44+ memory T cells can persist within the tumor site for extended periods of time post-treatment and may be important in achieving long term disease control. Anti-macrophage treatment might be especially efficacious in certain claudin-low BC patients as it could abolish immunosuppressive T12 macrophages and perhaps unleash B cell-mediated immunity. Further genetic studies are required to establish a functional role for helper T cells and antigen presenting B cells in these models.

Future studies using these models may help elucidate the mechanisms by which tumors of the same subtype differentially respond to treatment. A higher proportion of treatment-resistant circulating stem cells may explain the failure of CD8+ memory T cells to mount a strong anti-tumor response. Further studies are also required to elucidate the mechanisms that account for the phenotypic switch following the initial recruitment of macrophages to pathways that recruit immunosuppressive neutrophils. A more detailed analysis of the tumors that recur in the primary and metastatic sites is also needed. These preclinical studies will need to be extended to longitudinal studies in patients with new therapeutic combinations that can target potential immunological transitions to better improve long term progression free survival. Although Pexidartinib recently gained FDA approval to treat tenosynovial giant cell tumors (TNGCT)(13), its utility in other cancers has been limited because of both off target effects inhibiting c-kit and fms-like tyrosine kinase 3 (FLT3) and rare liver toxicity(39). Thus, Pexidartinib was discontinued for safety concerns in the I-SPY2 randomized clinical trial after accrual of only nine patients (40). One potential advantage of using preclinical models is to improve therapeutic regimens that very often lead to serious adverse events in patients that might be irreversible and cause long-lasting damage (15). Accordingly, we used a reduced (3.3 X lower dose) of PXB in combination with CTX to reduce potential liver toxicity. The low-dose combination treatment also yielded a significant survival benefit for primary tumors as compared to single agent treated T12 tumors. This illustrates the potential efficacy of drug combinations given at less than the maximum tolerated dose, but rather at a minimal effective dose. An alternative may be the use of Axatilimab, a novel mAB with high affinity for CSF-1R which is currently in Phase 2 trials for chronic graft-versus host disease (NCT04710576) without any reported liver toxicity. In addition, a number of novel macrophage inhibitors are in development. However, based upon the present studies it will be critical to test these in combination with the appropriate immunostimulatory chemotherapy.

Finally, the present study also further highlights the need to further understand the heterogeneity in immune cell subsets cell sub-populations. Future studies need to carefully examine their role in TNBC patients that do not respond to chemotherapy and to develop targeted treatments that can modulate B cell functionality like studies being done in TAMS. These results highlight the need to integrate newer techniques such as single cell RNA sequencing in combination with high dimensional imaging data to analyze patient samples in order to enhance our understanding of TILS and therapeutic response.

## Author Contribution

S.S., C.M.P., X.H.F.Z., and J.M.R. conceptualized the study, designed the experiments and wrote the manuscript. S.S. performed the experiments. C.H., S. A., D.A.P., Y.G., N.Z., and I.B., assisted with the mouse experiments and acquisition of data. L.C., Y.X., and S.L performed IMC staining and analysis. S.S., N.L., I.B., Y.W.W., Y.S., J.S., Z.L., and D.H. analyzed the bioinformatics data and discussed the project direction. S.S.,and C.M.P. analyzed clinical trial data.

## Acknowledgements

We would like to acknowledge Dr. Xi Chen for his help reviewing the manuscript, and Ipshita Thakur for illustrating the mice used in the treatment schemes. We would like to thank Jonathan Shepherd for his help with bioinformatic analysis. These studies were supported by grants NCI-CA148761(C.M.P and J.M.R.), Breast Cancer Research Foundation, NCI CA151293, US Department of Defense DAMD W81XWH-16-1-0073 and W81XWH-18-1-0574, Susan G. Komen CCR14298445, and McNair Medical Institute (X. H.-F. Z.) CPRIT grant RP160283(S.S.), the Dan L. Duncan Comprehensive Cancer Center (NCI: P30: CA125123) and Pathology Core of the Lester and Sue Smith Breast Center and T32CA203690-01A1 (D.A.P and Y.G). This project also was supported by the Advanced Technology Cores at BCM: Genomic and RNA Profiling Core (NCI: CA125123), Single Cell Genomics Core, Cytometry and Cell Sorting Core (CPRIT-RP180672), and Integrated Microscopy Core (NIH: DK56338 and CA125123; CPRIT: RP150578 and RP170719).

