## Supplementary material for "Chemotherapy coupled to macrophage inhibition leads to enhanced T and B cell infiltration and durable triple negative breast cancer regression": Supplmental Figures and Legends

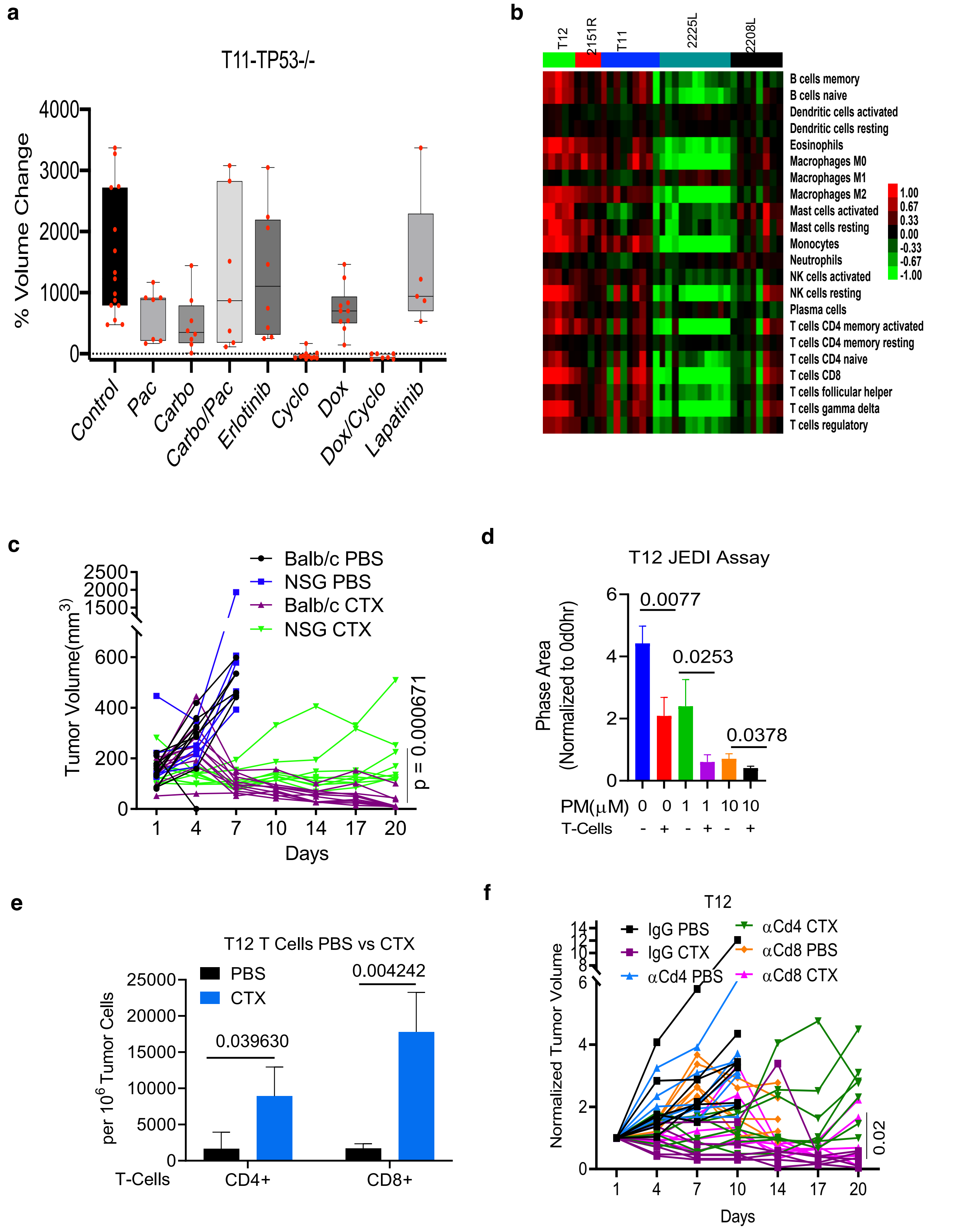


Supplementary Figure 1 for Figure 1, Singh et al.

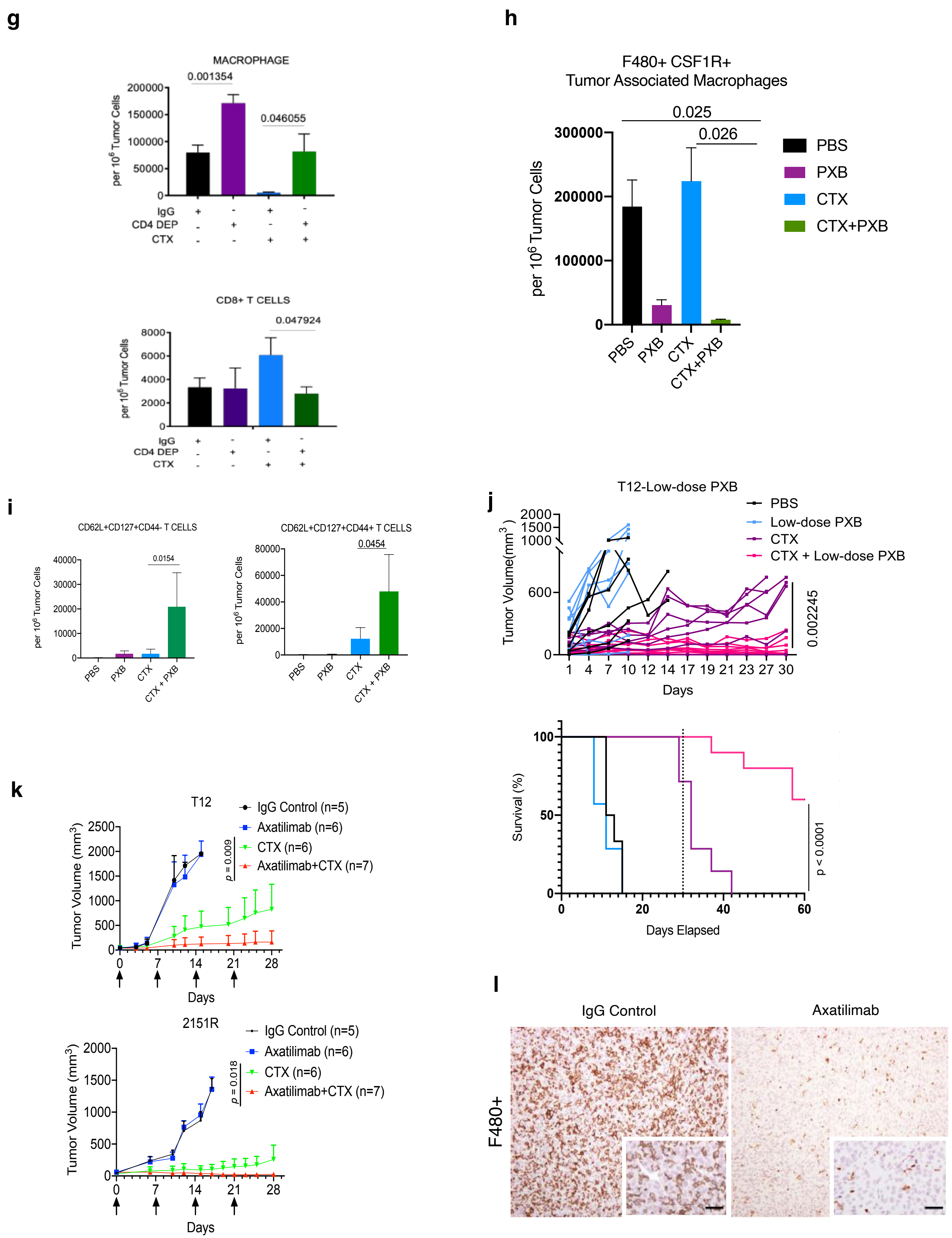


Supplementary Figure 1-2 for Figure 1, Singh et al.

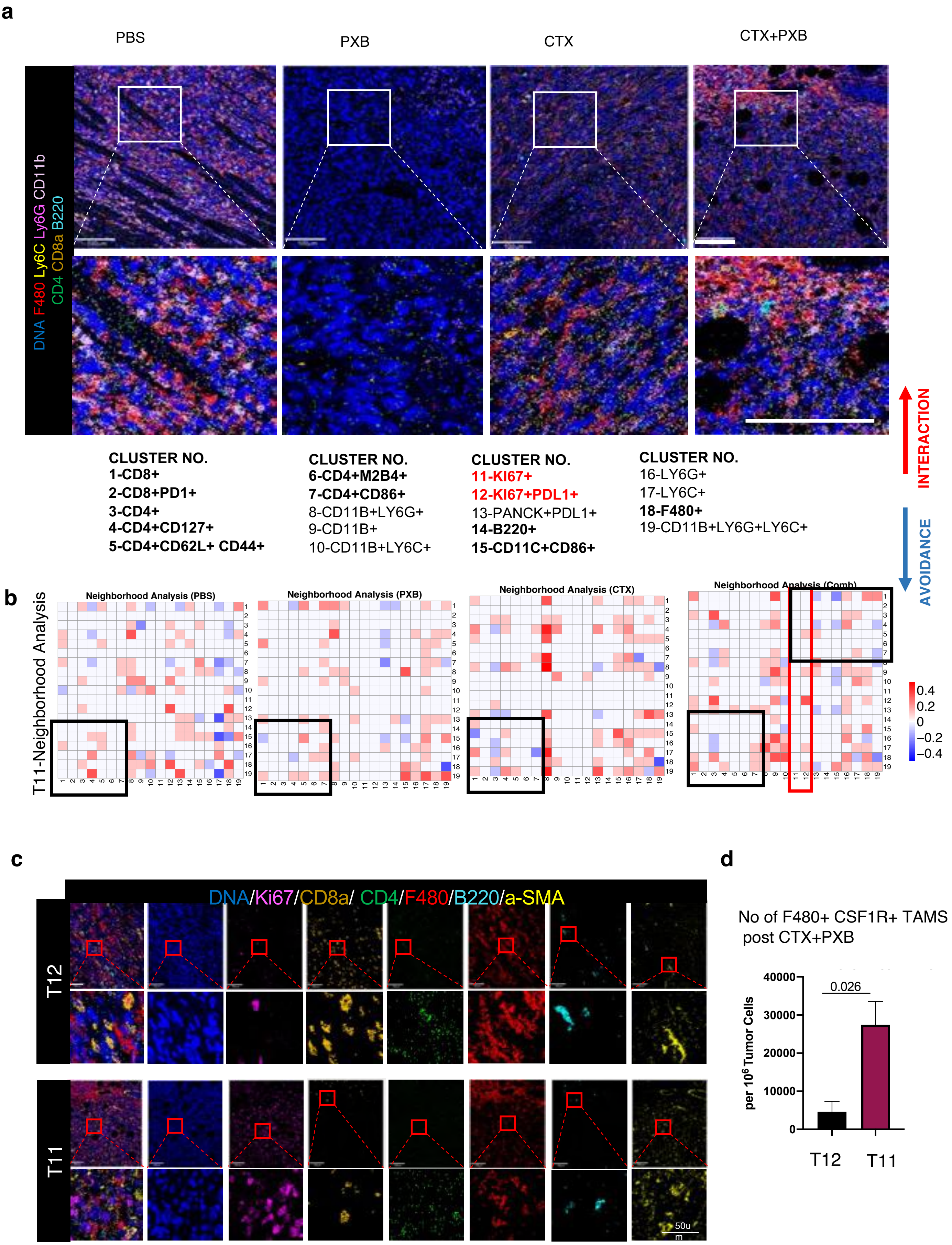


Supplementary Figure 2-1 for Figure 2, Singh et al.

**e**

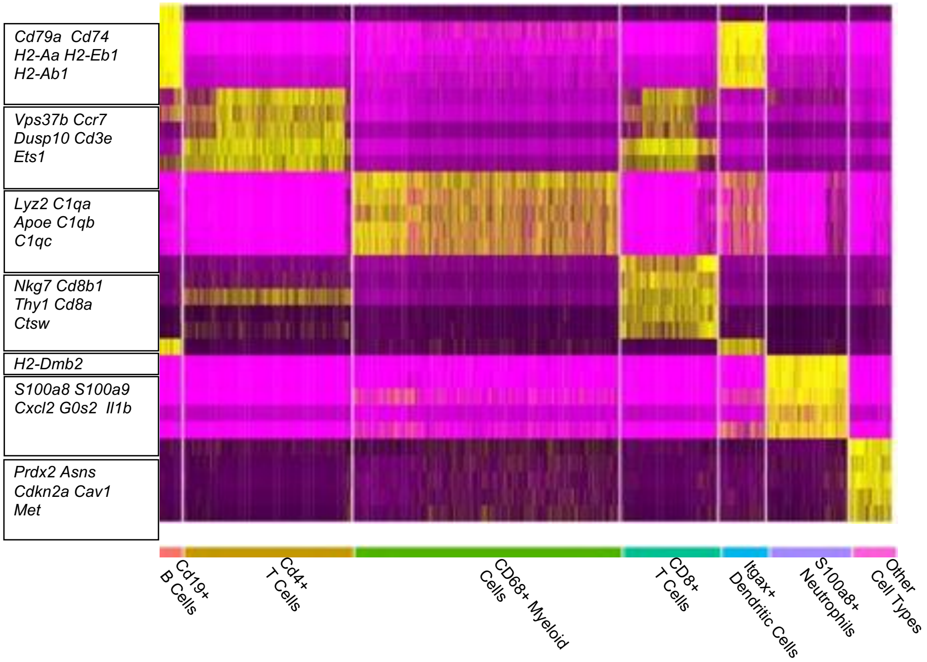


**f**

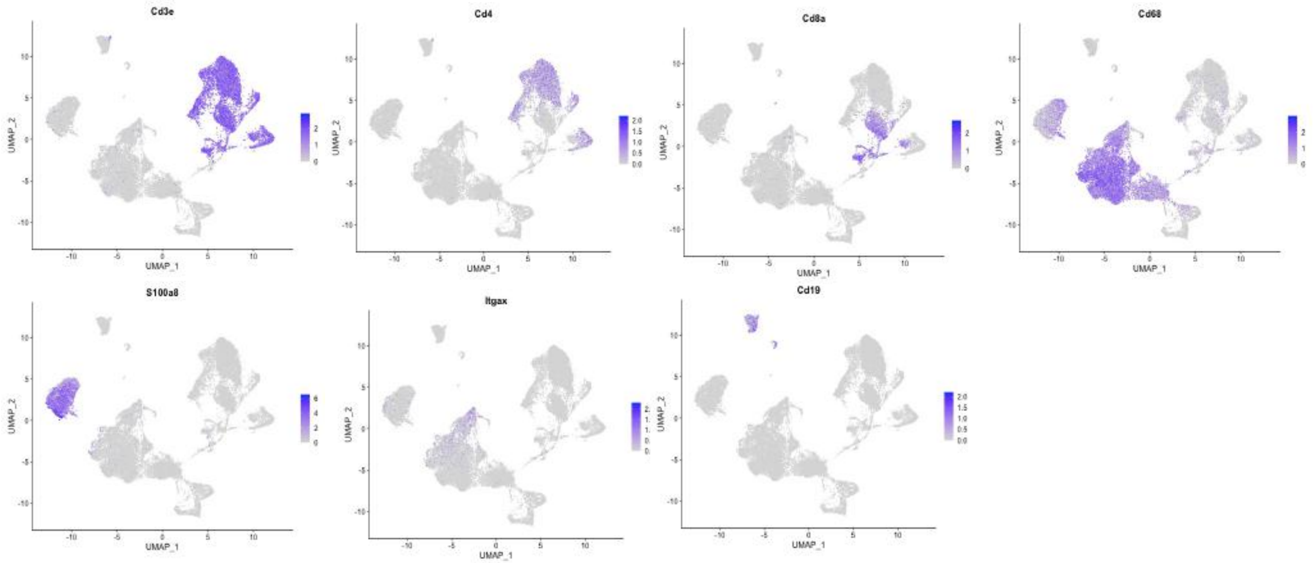


**g**

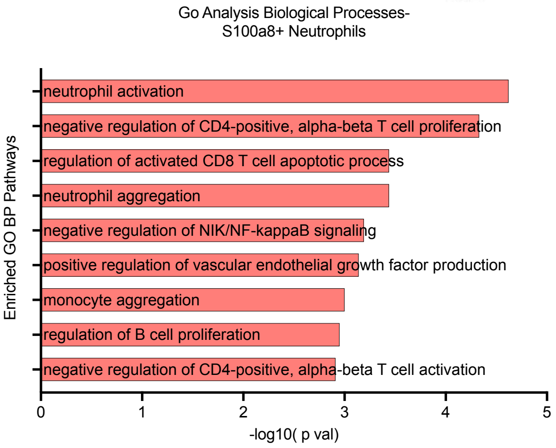


**h**

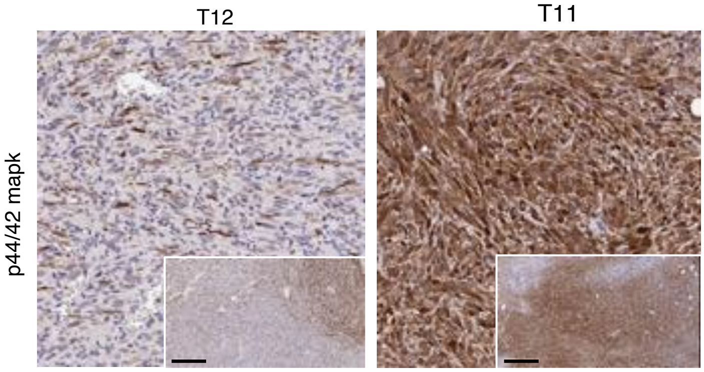


Supplementary Figure 2-2 for Figure 2, Singh et al.

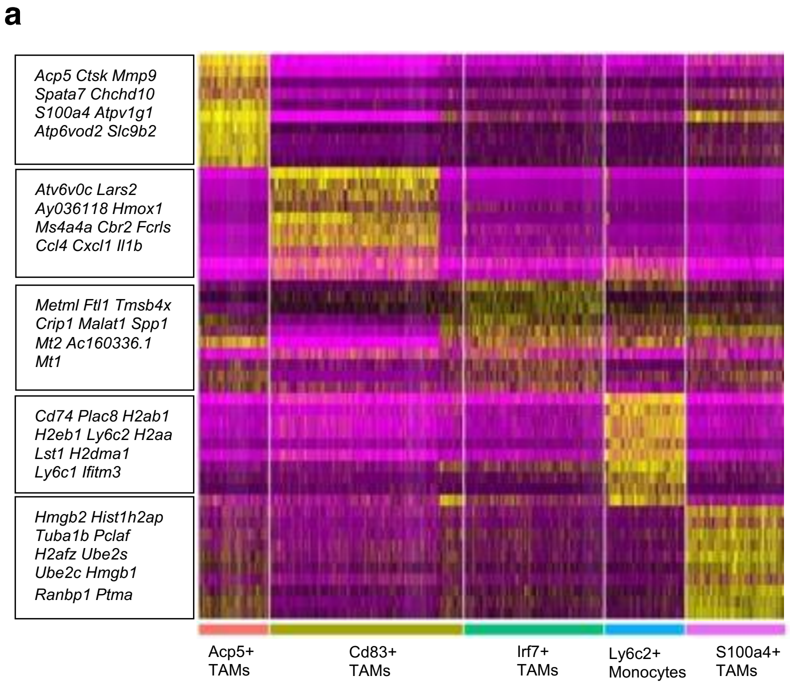


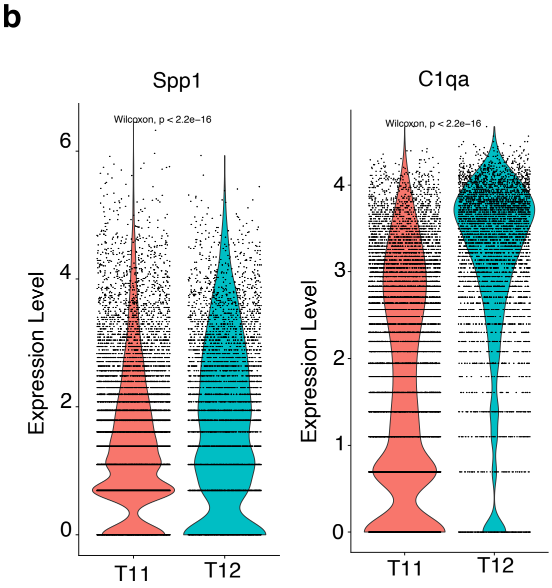


Pathways Analysis
T12 Cd68+ Myeloid Cells

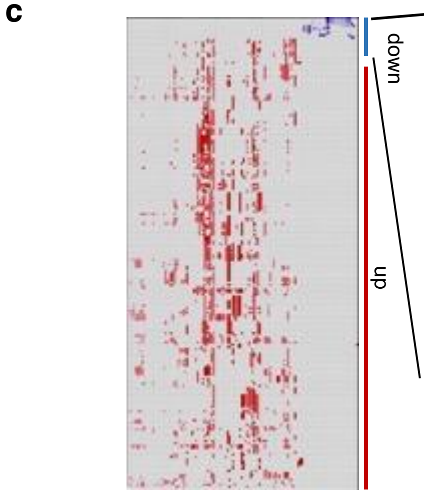


| NAME | NES | NOM p-val |
|---|---|---|
| Oxidative phosphorylation | -2.3721688 | 0 |
| Generation of precursor metabolites and energy | -2.0811594 | 0 |
| Mitochondrial organization | -1.7881126 | 0.00763359 |
| GOBP_EPITHELIUM_DEVELOPMENT | -1.2340722 | 0.1827957 |
| ATP metabolic processes | -1.2040108 | 0.16666667 |
| Protein containing complex subunit organization | -0.9362941 | 0.5684931 |
| Intracellular transport signaling | -0.9282206 | 0.60227275 |
| Membrane organization | -0.7036935 | 0.9078014 |
| Protein localization to organelles | -0.6916076 | 0.9444444 |

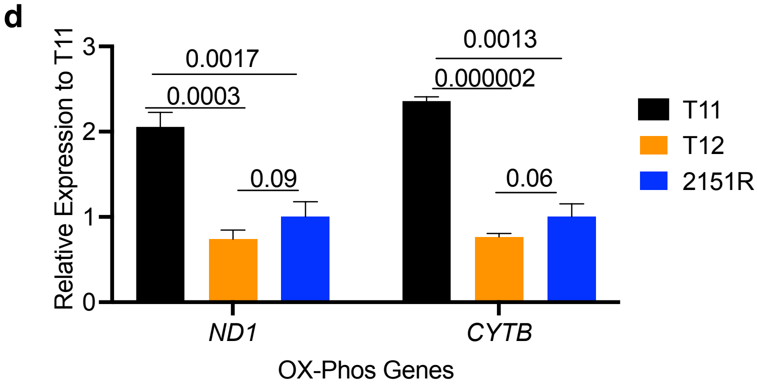


Supplementary Figure 3 for Figure 3, Singh et al.

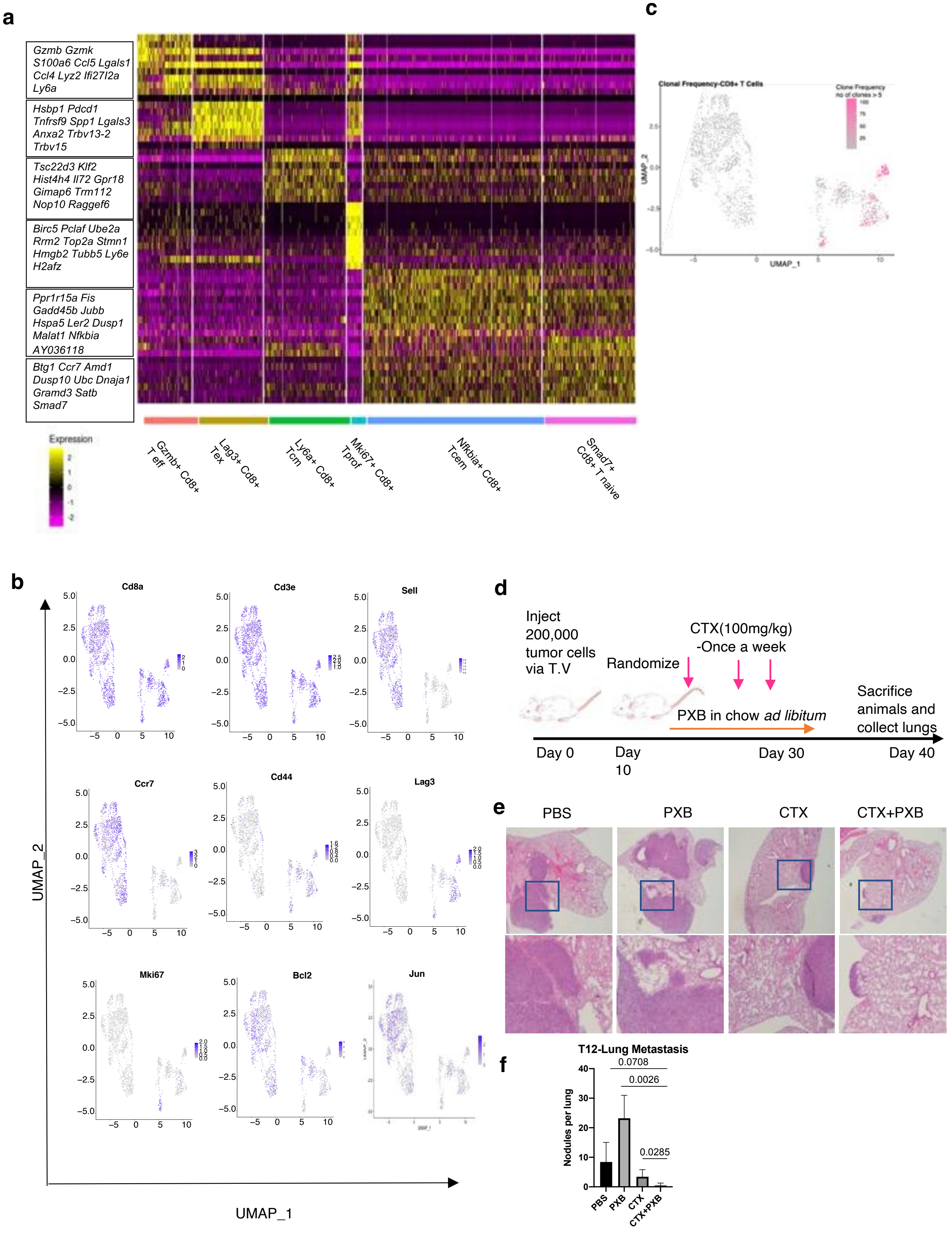


Supplementary Figure 4 for Figure 4, Singh et al.

g

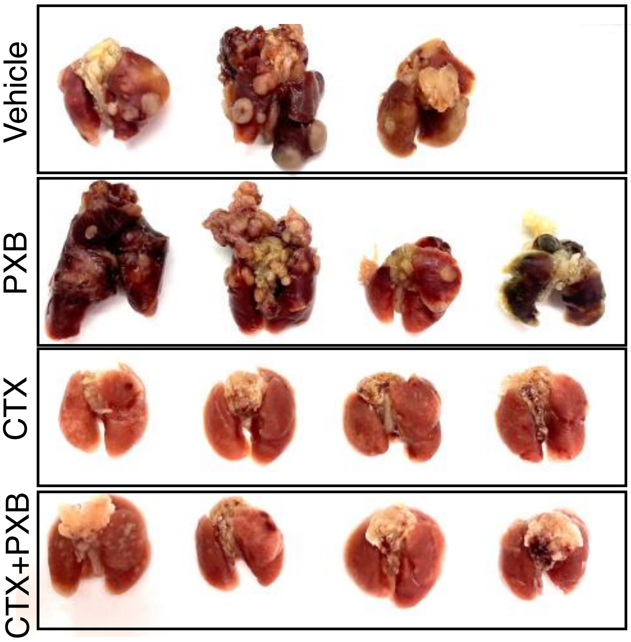


h

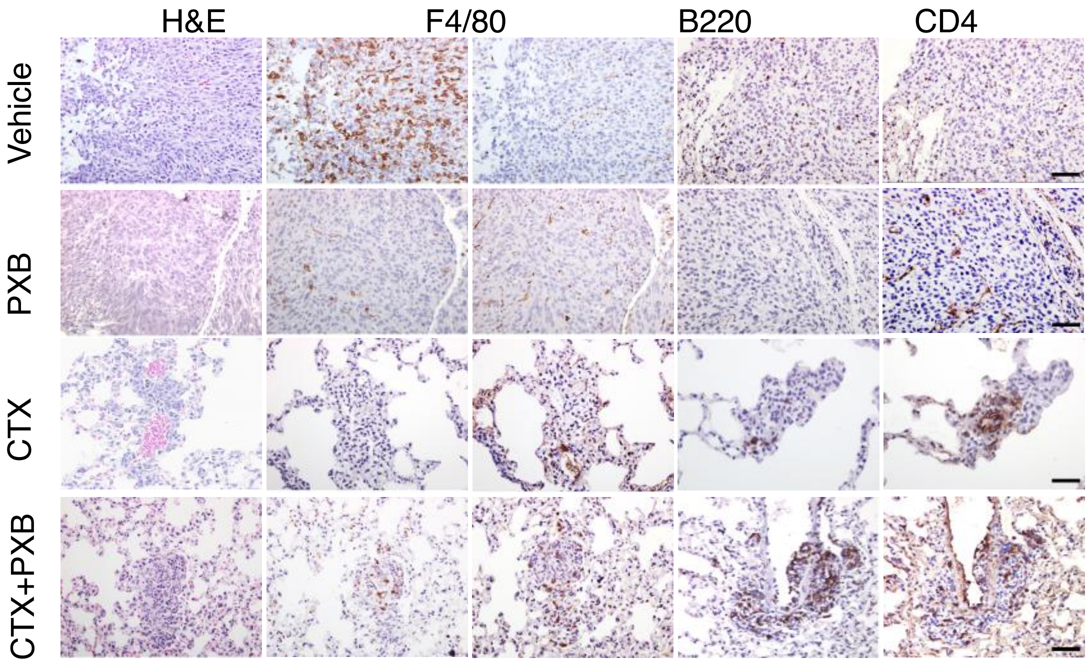


i

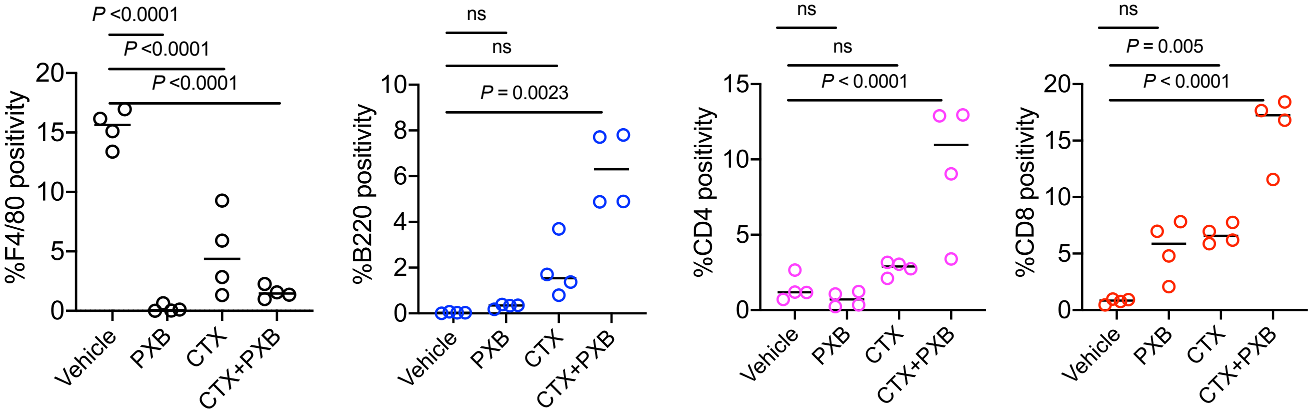


Supplementary Figure 4-2 for Figure 4, Singh et al.

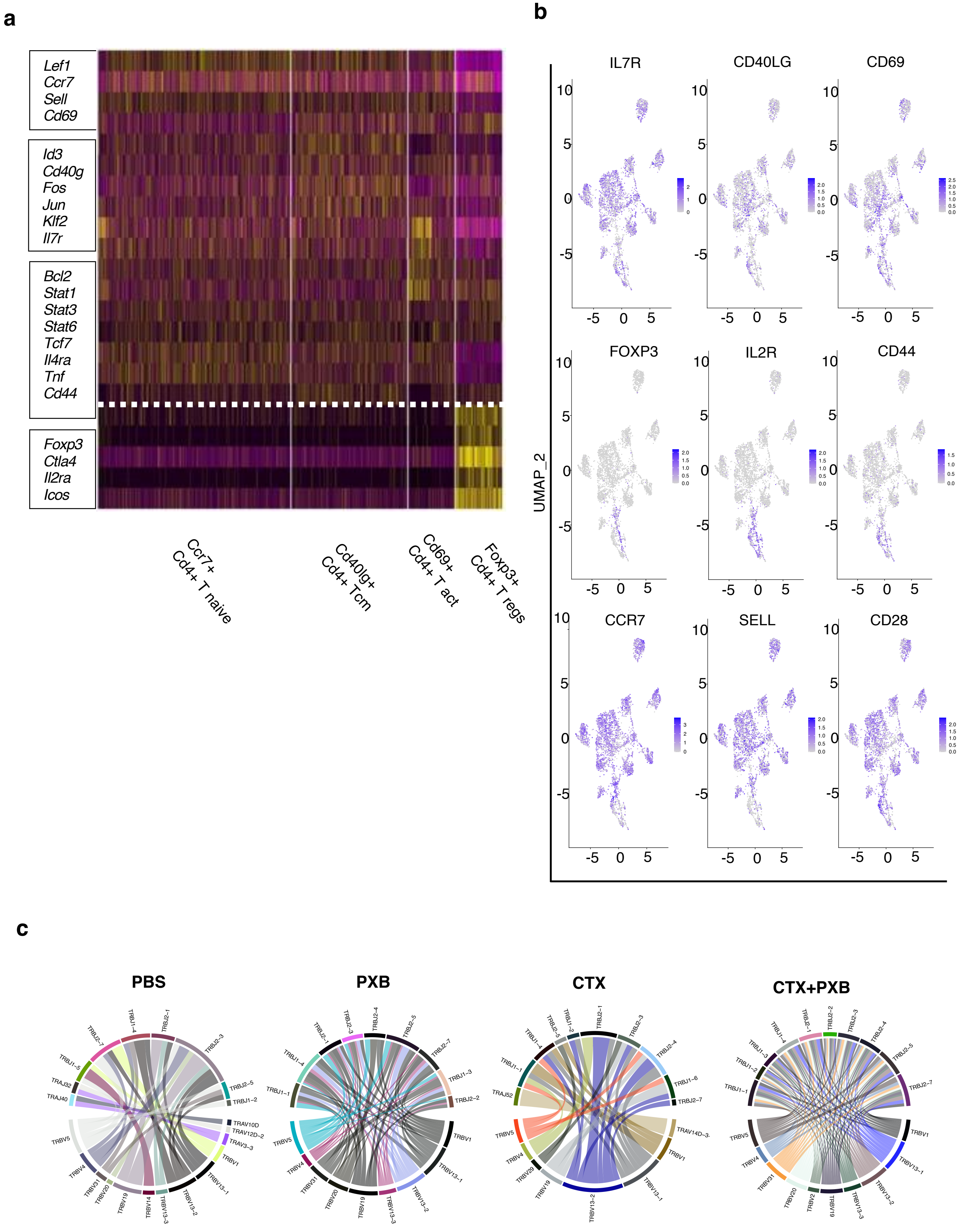


Supplementary Figure 5 for Figure 5, Singh et al.

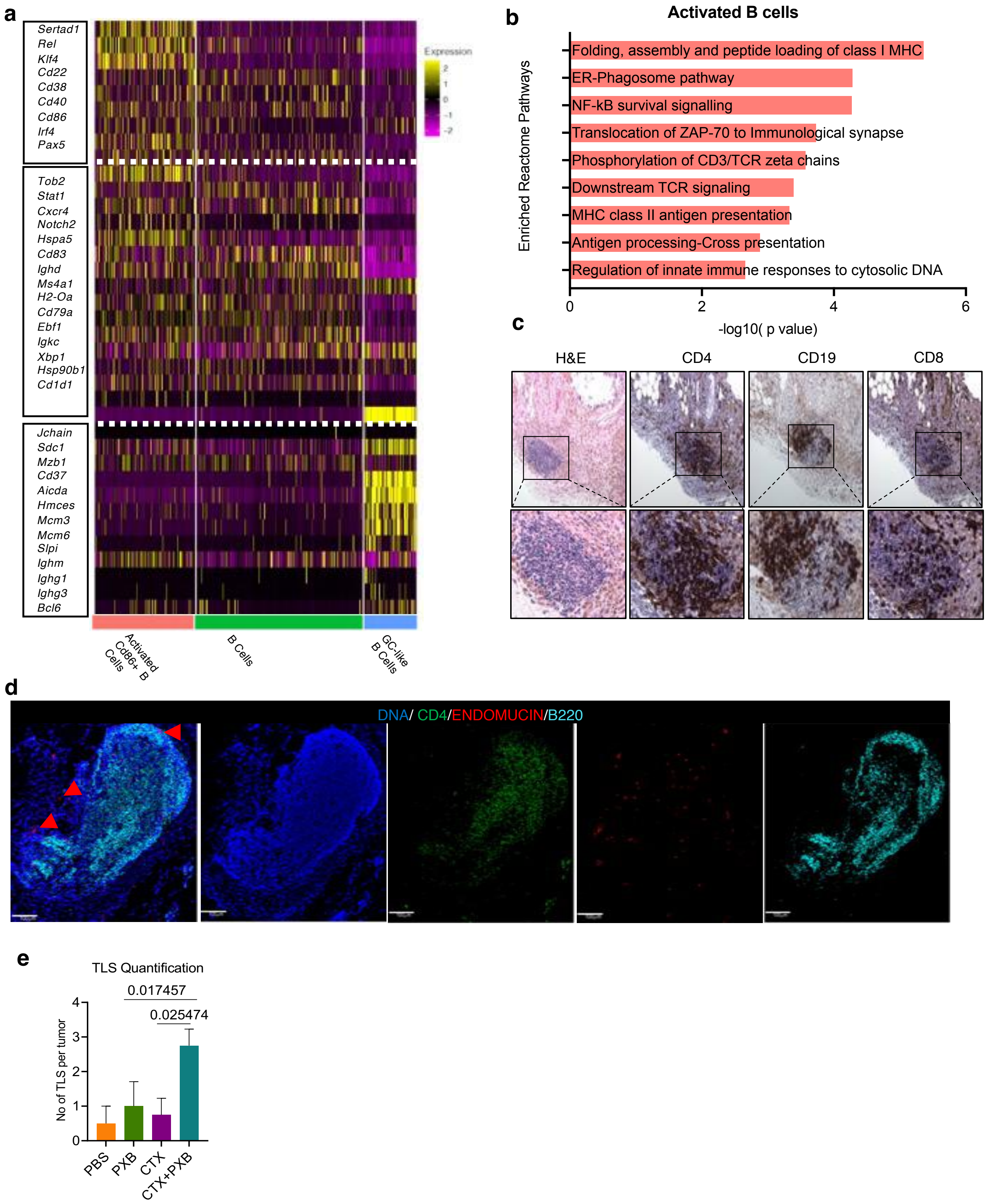


Supplementary Figure 6 for Figure 6, Singh et al.

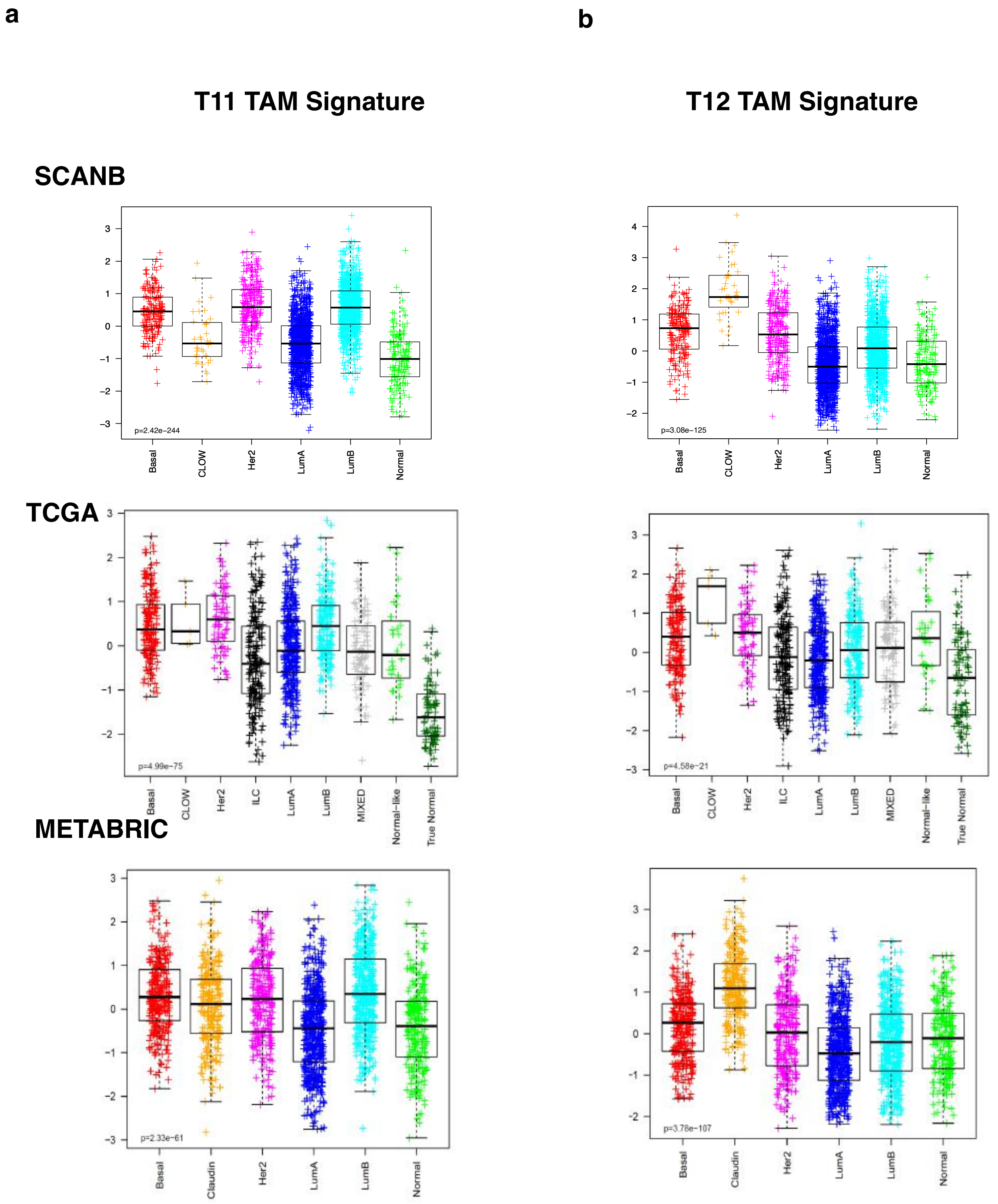


Supplementary Figure 7 for Figure 7, Singh et al.

### T Cell Gating

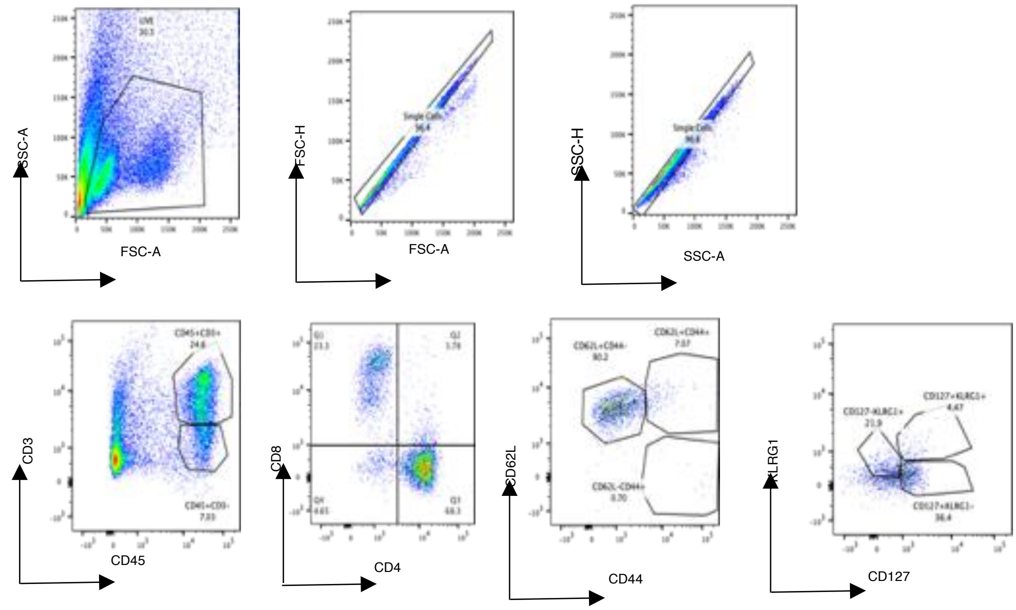


Supplementary Figure 8, Singh et al.

### Myeloid Cell Gating

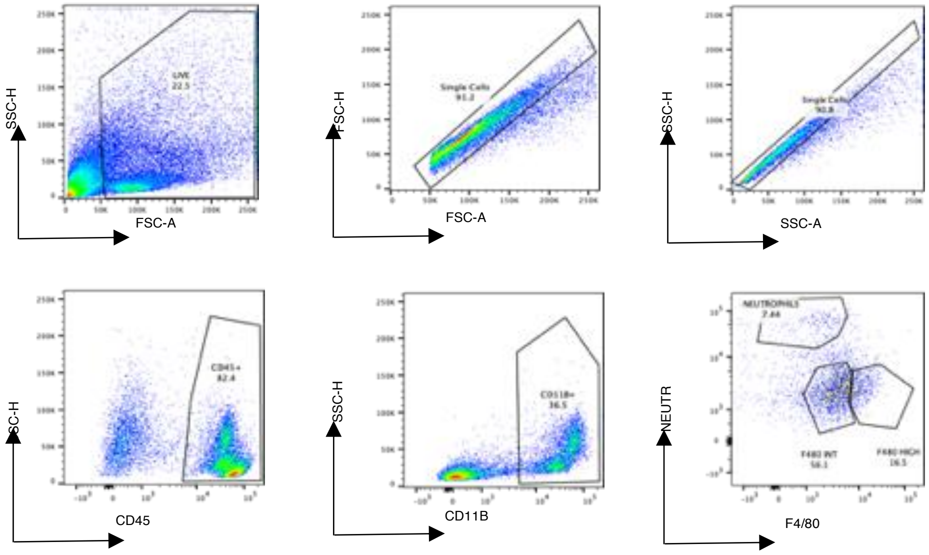


Supplementary Figure 9, Singh et al.

#### Supplemental Figure Legends

**Supplemental Figure 1. p53 -/- syngeneic TNBC GEM recapitulate human TNBC subtypes and respond to low dose immunostimulatory chemotherapy treatment.**
**a,** Treatment response using standard of care drugs on T11 mammary tumors. **b,** Cibersort analysis of murine TNBC models. **c**, Single agent CTX treatment led to a reduced tumor burden in Balb/c mice as compared to NSG mice. Data are shown as mean +/-S.E.M, n = 10 per group except NSG PBS, n = 5. Multiple unpaired t tests were used to determine significance. **d,** Co-culture of GFP+ T12 tumor cells and GFP targeting T cells with or without Phosphoramide Mustard (PM). Each treatment was performed in triplicates and repeated twice. **e,** Flow cytometry analysis of CD8+ and CD4+ T cells before and after low-dose CTX treatment, n = 6-7 mice per group. **f**, Antibody depletion of CD4+ T Cells leads to a significant decrease in response to CTX and increased tumor volumes while differences in response due to depletion of CD8+ T Cells neared statistical significance at day 17, p = 0.06 n = 5-7 mice per group. Multiple unpaired t tests were used to determine significance. **g,** Quantification of TAMs upon antibody depletion of CD4+ T cells after treatment and quantification of CD8+ T cells after CD4+ T cell depletion after treatment by single agent CTX**. h,** Quantification of F480+CSF1R+ TAMs after combination treatment with CTX and Pexidartinib using flow cytometry in T12 tumors, n= 5, significance was determined by unpaired t-tests. **i,** Quantification of CD8+ T memory cell subsets before and after treatment, n = 4-5 mice per group. Unpaired t-test was used to determine significance. **j,** Reduced tumor volume and improved survival in T12 mammary tumor bearing mice after treatment with CTX and 3.3X lower PXB(75 ppm). Number in parentheses show the specific n values of biologically independent mice per treatment group **k,** Combination of low-dose CTX with Axatilimab (mAB targeting CSF1R) leads to a reduced primary tumor burden of T12 and 2151R p53-/- GEMMs following 4 weekly treatments (arrows). **l,** Immunohistochemistry for F4/80+ TAMs following 1 dose of IgG or Axatilimab treatment.

**Supplemental Figure 2. Combination therapy leads to an expansion of CD8+/CD4+ T cells and B cells in responsive T12 tumors. a,** Imaging mass cytometry analysis of the tumor immune microenvironment in T11 tumors before and after treatment. Representative images overlaid with 7 markers (F480, Ly6C, Ly6G, CD11b, CD4, CD8a, B220) for each treatment group. **b**, Neighborhood analysis of T11 tumors in which the color of the squares indicates significant pairwise interactions or avoidance between PhenoGraph defined cellular metaclusters. Highlighted interactions include CD8+/CD4+ T cells (clusters 1-7), B220+ B cells(cluster-15) and CD11C+ CD86+ Dendritic cells (cluster 16). Three Regions of Interest (ROI) were ablated per tumor section. n= 3-5 independent biological replicates per treatment group. **c**, Representative IMC images of T12 and T11 tumors after combination therapy highlighting the spatial localization of CD8+/CD4+ T cells and B cells with respect to proliferating KI67+ cells and F480+ TAMs. **d,** Flow cytometry quantification of F480+ CSF1R+ T12 and T11 tumor associated macrophages after combination therapy, n=3 independent biological replicates per group. **e,** Gene expression heatmap for CD45+ immune cells before and after treatment for all treatment groups. **f** Feature plots showing the expression levels of murine immune cell marker genes-*Cd3e, Cd4, Cd8a, Cd19, Cd68, S100a8, Itgax,* in all 4 treatment groups. **g,** GOBP analysis of *S100a8*+ neutrophils that increase in T12 tumors after single agent PXB treatment. **h,** IHC showing elevated pMAPK signaling in T11 tumors pre-treatment

**Supplemental Figure 3. TAMs in responsive (T12) and non-responsive (T11) tumors are phenotypically distinct. a,** Heatmap showing top 10 upregulated genes per TAM sub-cluster for T12 and T11. **b,** Quantification of additional TAM marker genes. **c,** Heatmap representing top upregulated and downregulated metabolic pathways in T12 TAMs. **d,** qPCR analysis of T12, T11 and 2151R tumors using select genes related to oxidative phosphorylation (n=3 independent biological replicates per tumor model, p value calculated using two-sided T test)**.**

**Supplemental Figure 4. Combination therapy leads to polyclonal expansion T cells that exhibit memory cell phenotypes. a,** Gene expression heatmap top genes for each *Cd8*+ T cell sub-cluster **b,** Feature plots showing the expression of select memory/exhaustion/activation markers in *Cd8*+ T cells. **c.** V(D)J analysis of *Cd8*+ T cells for all 4 treatment groups showing polyclonal expansion of *Cd8*+ T cells after combination treatment. **d,** Treatment schema depicting treatment for mice bearing T12 lung metastasis. **e, f,** Representative images and quantification of surface metastatic lung nodules in all 4 treatment groups-PBS, PXB, CTX, CTX+PXB, n= 5-6 animals per group, $p < 0.05$).
**d,** Schema for T12 rechallenge experiments and treatment. **e,f,** Quantification of mice that completely or partially (tumor volume < 50 % of control tumors) rejected T12 mammary tumors injected into the contralateral mammary gland of previously treated T12 tumor bearing mice, n=10. **g,** Images of visible lung metastatic nodules following Veh, PXB, CTX and CTX+PXB. **h,** Immunohistochemistry of T12 macro and micro-metastatic lung lesions for H&E, F480+ TAMs, B220+ B- and CD4/8+ T-cell markers following 30-day treatment of GEMMs**. i,** Quantification of immunohistochemistry showing % positivity of F480, B220, CD4 and CD8 following Veh, PXB, CTX and CTX+PXB.

**Supplemental Figure 5. Cd4+ T cell and B cell play an important role in mediating long term tumor regression. a,** Heatmap showing top upregulated genes per *Cd4*+ T cell sub-cluster. **b,** Feature plots showing the expression of select memory/exhaustion/activation markers in CD4+ T cells across all 4 treatment groups. **c,** Clonal frequency of Cd4+ T cells in all 4 treatment groups and chord diagrams representing unique V-J region pairings in *Cd4*+ T cells in T12 tumors before and after treatment.

**Supplemental Figure 6. Activated B cells expand after combination therapy and may be the main APCs within the tertiary lymphoid structures. a,** Heatmap showing highly expressed genes in every B cell sub-cluster**. b,** REACTOME analysis of *Cd86*+ Activated B cells(ABCs) expanded after combination treatment. **c,** H&E and IHC images of TLS using T cell (CD4+ or CD8+) and B cell (CD19+) markers. **d,** Representative imaging mass cytometry images as well as single channel images depicting the localization of HEV marker Endomucin with B220+ B cells and CD4+ T cells withing TLS. **e,** Quantification of TLS in all 4 treatment groups in T12 tumors ($p < 0.05$). P value was computed using two-sided T test.

**Supplemental Figure 7. T12 TAM signature is upregulated in patients with Claudin-low breast cancer in all and TNBC only patient samples. a, Downregulation** of T11 TAM signature in all claudin-low breast cancer patients in SCANB, TCGA and METABRIC clinical trial datasets. **b,** Upregulation of T12 TAM signature in all claudin-low breast cancer patients in SCANB, TCGA and METABRIC clinical trial datasets.

**Supplemental Figure 8. T cell gating strategy.**

**Supplemental Figure 9. Myeloid cell gating strategy.**
